# Selective cholinergic stimulation of the medial septum-diagonal band of Broca via DREADDs improves spatial learning in healthy rats

**DOI:** 10.1101/2022.08.02.502516

**Authors:** Stephan Missault, Sam De Waegenaere, Lauren Kosten, Annemie Van der Linden, Marleen Verhoye, Georgios A. Keliris

## Abstract

The septohippocampal pathway plays an important role in learning and memory. It projects from the medial septum-vertical limb of the diagonal band of Broca (MSDB) to the hippocampus and provides the latter with its main cholinergic innervation. To assess the importance of cholinergic selectivity and timing of MSDB stimulation in modulating learning and memory, we directly compared the effects of several MSDB stimulation strategies in healthy rats. We evaluated the effects of DREADD-mediated selective cholinergic neuronal MSDB stimulation and nonselective neuronal MSDB stimulation on spatial learning and memory in the appetitive radial arm maze and on resting-state brain networks using resting-state functional MRI. DREADDs were activated with the novel DREADD agonist J60. Selective cholinergic MSDB stimulation during – but not after – radial arm maze training improved spatial learning compared with J60-treated sham rats and had no effect on working memory or reversal learning. J60-treated sham rats had a worse working memory than saline-treated sham rats during the reversal phase of the radial arm maze task, suggesting an adverse effect of chronic use of J60. Nonselective MSDB stimulation during training resulted in a loss of appetite and exclusion from the radial arm maze training. Acute selective cholinergic and nonselective MSDB stimulation induced decreased functional connectivity (FC) in the default mode-like network. In addition, acute nonselective MSDB stimulation resulted in increased intrahippocampal FC, while selective cholinergic MSDB stimulation led to globally increased FC with the nucleus accumbens. While the combined effect of radial arm maze learning and the necessary chronic food restriction with or without chronic MSDB stimulation had no observable effect on resting-state networks, chronic food restriction alone globally increased FC in the brain.

## Introduction

The septohippocampal pathway consists of cholinergic, GABAergic and glutamatergic neurons projecting from the medial septum and the vertical limb of the diagonal band of Broca (MSDB) in the forebrain to the hippocampal formation and plays an important role in learning and memory^1^. MSDB lesions have been shown to result in impaired learning and memory in hippocampus-dependent tasks in rodents^2^. A major role herein has been attributed to the cholinergic projection neurons, which constitute approximately 60% of the septohippocampal neurons in the rat^3^ and provide the hippocampus with its main cholinergic innervation^4^. The importance of acetylcholine in cognition was underlined by early observations of a selective loss of basal forebrain cholinergic neurons in Alzheimer’s disease^5, 6^ and the cognitive benefits, though modest, of acetylcholinesterase inhibitors in patients with Alzheimer’s disease^7^. A disadvantage of these drugs, however, is that they have common adverse effects^8^, which could be avoided by limiting the acetylcholine increase to the cognition-relevant pathways in the brain. This could be achieved by directly stimulating the cognition-relevant cholinergic nuclei in the basal forebrain.

Stimulation of the MSDB has been shown to modulate learning and memory in rodents: both improvement and impairment have been observed. The effect of MSDB stimulation on cognition seems to be dependent on the applied method, cell-type specificity, timing of stimulation, frequency of stimulation, developmental phase, and the absence or presence of pre-existing cognitive deficits. Several techniques have been employed to stimulate the MSDB, including electrical stimulation/deep brain stimulation, pharmacological manipulation through intraseptal injection of different receptor (ant)agonists, optogenetics and chemogenetics. In most behavioural studies, spatial learning and memory tasks were employed to assess the cognitive effect of MSDB stimulation. Depending on the task, different forms of memory can be investigated: reference memory/acquisition learning, reversal learning and working memory. Using the example of the radial arm maze, reference memory, a form of long-term memory, refers to remembering the fixed location of the rewards relative to surrounding visual cues. Learning the fixed location of the rewards for the first time is referred to as acquisition learning. After some time, the location of the rewards can be changed after which animals have to learn the new fixed location of the rewards. This process is called reversal learning and is used to test cognitive flexibility. Working memory, a form of short-term memory, refers to remembering which arms of the maze were previously visited within a trial, independent of the learning phase (acquisition/reversal).

Most studies that reported cognitive improvement after MSDB stimulation only observed this in animal models with cognitive deficits. Deep brain stimulation of the MSDB resulted in improved spatial learning (reference memory) in rats after traumatic brain injury^9, 10^, improved spatial reference memory retention in a cholinergic lesion rat model of dementia^11, 12^, improved spatial learning (reference memory) in a rat model of temporal lobe epilepsy^13, 14^ and improved novel object recognition in a mouse model of temporal lobe epilepsy^15^. Activation of 5-HT 2A receptors and both activation and blockade of 5-HT 6 receptors in the MSDB improved spatial working memory in hemiparkinsonian rats^16, 17^. Intraseptal administration of the cholinergic receptor agonist carbachol facilitated encoding of new information in aged rats^18^ and the muscarinic agonist oxotremorine improved spatial working memory in aged rats^19^. Selective optogenetic stimulation of parvalbumin-positive GABAergic MSDB neurons improved object location memory in a mouse model of Alzheimer’s disease when stimulated at 40 Hz but not at 80 Hz during the retrieval phase of the task^20^. MSDB stimulation in the control animals in these studies – if studied – most often had no effect on cognition^9, 13, 14, 19^, sometimes an adverse effect on cognition^10, 18^ and other times a beneficial effect on learning and memory^10, 16, 17^. In one study, deep brain stimulation of the MSDB with theta frequency (7.7 Hz) caused sham rats to use fewer random strategies in a spatial learning (reference memory) task, while gamma frequency stimulation (100 Hz) in the sham rats caused a deficit in this task^10^. This outcome was not reproduced in another study from the same research group in which MSDB deep brain stimulation at theta frequency had no effect on the spatial learning task in sham animals^14^. Activation of 5-HT 2A receptors or 5-HT 6 receptors in the MSDB was found to enhance spatial working memory in sham rats^16, 17^. A few other studies have observed improved cognition upon MSDB stimulation in healthy animals. Deep brain stimulation improved the retention of a brightness-discrimination task in rats^21^, facilitated the acquisition of a visual light/dark discrimination task in rats^22^ and improved retention of a lever-press operant conditioning task with food reward in mice^23^. Post-training intraseptal administration of glutamate, the muscarinic acetylcholine receptor agonist oxotremorine, norepinephrine or the GABAA receptor antagonists picrotoxin or flumazenil facilitated retention in a passive avoidance task in rats^24, 25^. Pre-training intraseptal administration of acetylcholinesterase inhibitor physostigmine or GABAA receptor antagonist bicuculline improved memory acquisition in a passive avoidance task in rats^26^. Activation of 5-HT 1A receptors facilitated acquisition of a spatial discrimination task in mice^27^. Nonselective neuronal optogenetic stimulation improved spatial working memory in rats^28^ and optogenetic stimulation of CA1-projecting septohippocampal neurons improved contextual fear memory retrieval in mice^29^.

Most of the MSDB stimulation strategies that were able to improve memory were not selective for a specific neuronal subpopulation. With deep brain stimulation several neuronal subpopulations are stimulated and also in the case of pharmacological manipulation different neuronal subpopulations are usually involved since the targeted receptors are often expressed on multiple neuronal subpopulations. 5-HT1A and 5-HT 2A receptors for instance have been shown to be present on both parvalbumin-positive GABAergic neurons and cholinergic neurons in the MSDB of rats^30, 31^. From the above-mentioned studies it cannot be deduced whether selective stimulation of cholinergic MSDB neurons could have a similar or differential effect on cognition compared with nonselective stimulation. Optogenetic and chemogenetic techniques can be employed to target specific neuronal subpopulations. Though optogenetic and chemogenetic techniques have already been employed to activate or inactivate a specific neuronal subpopulation in the MSDB or to (in)activate MSDB neurons nonselectively, with various effects on cognition, no studies so far have directly compared the effects of a nonselective and a selective MSDB neuronal stimulation approach on cognition.

In our study, we aimed to compare the effects of selective stimulation of cholinergic MSDB neurons and nonselective neuronal MSDB stimulation using chemogenetics on spatial learning and memory in healthy rats. We focused on cholinergic neurons given the prominent role of acetylcholine in cognition. While the above-mentioned studies typically focused only on one form of memory, we evaluate the effect of our MSDB stimulation strategies on spatial acquisition learning/reference memory, reversal learning, and spatial working memory. We opted for a chemogenetic technique (Designer Receptors Exclusively Activated by Designer Drugs or DREADDs) rather than optogenetics since chemogenetics have more translational value as a potential therapy in humans than optogenetics. While both techniques require neurosurgery, chemogenetic approaches do not require the implantation of any instrumentation as is the case for optogenetics and deep brain stimulation, which can give rise to hardware-related infections^32^, and neither is there any need for a replacement surgery of the neurostimulator in the long term.

After successful expression of DREADDs in the brain area of interest, they can be activated by systemic administration of a DREADD agonist. While clozapine N-oxide (CNO) has classically been used as agonist for the human muscarinic acetylcholine receptor-based DREADDs (including the excitatory hM3Dq DREADD used in this study), it has recently been shown that CNO cannot cross the blood-brain barrier and acts on DREADDs through reverse metabolism to its parent compound clozapine, an atypical antipsychotic that can bind to many different endogenous receptors and can produce numerous behavioural effects^33^. Moreover, it has been shown that CNO administration can produce behavioural effects in rats even in the absence of DREADDs^34^ and that CNO has clozapine-like behavioural effects at doses commonly used to activate DREADDs in both rats and mice^35^. These observations emphasise the need for a DREADD-free control group treated with the DREADD agonist when performing DREADD experiments^34, 35^. Moreover, individual variation in CNO-to-clozapine conversion and/or clozapine metabolism between subjects may lie at the basis of a differential response of animals to the same CNO dose (both in terms of the DREADD response and the undesired adverse effects), as indicated by widely varying plasma levels of clozapine in rats after administration of the same dose of CNO and a considerably variable behavioural response of animals to same doses of CNO^35^. Hence, novel high-potency DREADD agonists have been introduced by the research group of Michael Michaelides that can cross the blood-brain barrier and act directly on DREADDs: JHU37152 and JHU37160^36^. We used JHU37160 (“J60”) in this study to activate hM3Dq DREADDs. Very few studies have used J60 for the activation of DREADDs so far^36, 37, 38^. The first study from Michael Michaelides’ group tested several doses of J60 (ranging from 0.01 to 1 mg/kg) in mice and rats and did not observe adverse effects of J60 on locomotion and electrical stimulation evoked responses in DREADD-free rodents^36^. The second study used a dose of 0.01 mg/kg J60 in mice and did not demonstrate effects on electromyography of the genioglossus muscle, fluorodeoxyglucose uptake in the tongue and respiratory measures in DREADD-free animals^38^. In the final study, a dose of 0.1 mg/kg J60 was used in mice, but no DREADD-free animals were included to test for J60-related adverse effects^37^. None of these studies, however, used J60 in a chronic manner. Since we proposed to inject animals almost daily with J60 for a total of 14 days over a period of 19 days (five days J60 – two days no J60 – five days J60 – three days no J60 – four days J60), we deemed the inclusion of both a J60-treated DREADD-free control group and a saline-treated DREADD-free control group necessary to investigate possible J60-related adverse effects due to chronic/repeated J60 administration. Based on personal communication with Michael Michaelides, a dose of 0.1 mg/kg J60 was chosen for our rat study.

While acetylcholine is important for learning and memory, and enhancing acetylcholine levels can have beneficial effects on cognition, hippocampal acetylcholine levels need to be low during slow-wave sleep for an optimal memory consolidation^39^. Increasing acetylcholine levels during the first part of sleep, a period with a high percentage of slow-wave sleep, has been shown to completely block slow-wave sleep-related consolidation of declarative memories in human subjects^39^. However, a high cholinergic tone is observed during active wakefulness and rapid eye movement (REM) sleep and is likely needed for memory processes taking place during those stages, including memory encoding^40, 41^. Hence, we hypothesised that increasing acetylcholine levels during memory training (during active wakefulness, in the animals’ dark phase) through MSDB stimulation would enhance memory formation and improve spatial learning, while increasing acetylcholine during the first part of the animals’ resting phase (light phase) would interfere with memory consolidation during slow-wave sleep, resulting in impaired spatial learning.

Most MSDB studies that showed improved cognition initiated MSDB stimulation immediately before the behavioural task (acquisition of a task or working memory task) started, either by intraseptal injection or through electrical stimulation, and continued stimulation throughout the test (as long as the injected drug’s effects lasted or until the end of the test in case of deep brain stimulation)^10, 13, 14, 16, 17, 18, 19, 26, 27, 28^. Using a similar approach, we aimed to improve memory by initiating MSDB stimulation shortly before the behavioural testing starts.

Summarised, first of all we aimed to elucidate the importance of cholinergic specificity when stimulating the MSDB for memory improvement. To this purpose, we directly compared the effects of a selective cholinergic MSDB stimulation approach and a nonselective neuronal MSDB stimulation approach on different forms of learning and memory: spatial reference memory/acquisition learning, reversal learning, and working memory. Secondly, we wanted to confirm our hypothesis that cholinergic stimulation during memory training would improve memory and cholinergic stimulation after memory training would impair memory. Thirdly, we wanted to evaluate the chronic use of the novel DREADD agonist J60 by comparing J60-treated and saline-treated DREADD-free animals. Finally, we wanted to evaluate the effect of i) acute MSDB stimulation, and ii) chronic MSDB stimulation and learning on whole-brain resting-state networks using functional magnetic resonance imaging (fMRI). Learning-induced changes in resting-state networks have been described^42, 43^, but to our knowledge the effect of MSDB stimulation on resting-state networks has not been investigated so far.

## Material and Methods

### Animals

Eighty-two healthy adult male ChAT-Cre (n=46) and wild type (WT) (n=36) Long-Evans rats were used in this study. Rats were bred in-house using transgenic LE-Tg(Chat-Cre)5.1Deis rats^44^ purchased from the Rat Resource & Research Center P40OD011062 (RRRC, USA) and WT Crl:LE Long-Evans rats purchased from Charles River Laboratories (Italy). Animals were initially group-housed in a temperature- and humidity-controlled room on a 12-hour reversed light-dark cycle with standard food and water available *ad libitum*. Later on, animals were single-housed when food restriction was necessary. Water was always available *ad libitum*. Animals were treated in accordance with EU directive 2010/63/EU. Animal experiments were approved by the animal ethics committee of the University of Antwerp, Belgium (ECD 2018-58).

### Study design

There were two cohorts of animals in this study. The first and main cohort consisted of five groups that were subjected to stereotaxic surgery, memory training with food restriction required for the behavioural task, and repeated MRI (before and after memory training) (Fig.1A). The second cohort consisted of a group of animals without any surgery that were subjected to food restriction and repeated MRI, but without memory training (Fig.1B).

**Figure 1.**
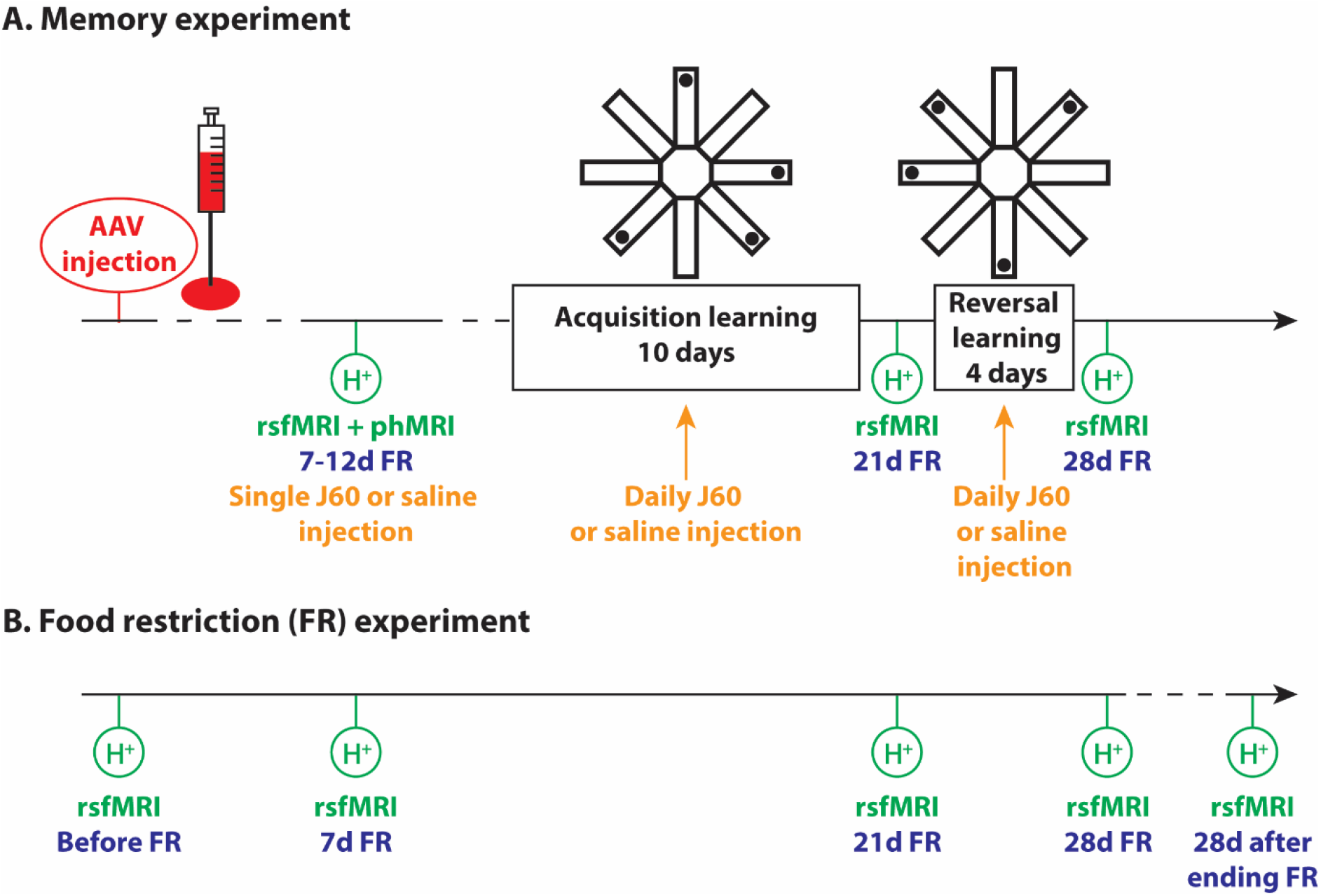
Study design. A. Memory experiment. Rats were subjected to intracranial adeno-associated viral (AAV) injections for the expression of DREADDs and/or mCherry in the medial septum-vertical limb of the diagonal band of Broca (MSDB). Minimally six weeks later, rats were subjected to combined resting-state functional MRI (rsfMRI) and pharmacological functional connectivity MRI (phMRI) with the novel DREADD agonist J(HU371)60 or the vehicle saline. For this scanning session, rats were food-restricted for 7-12 days. Next, rats were subjected to memory training in the radial arm maze. At first, rats had to memorise the fixed location of four rewards for 10 days during the acquisition learning phase. Rats received daily injections of J60 or saline during this learning phase. Next, rats were again subjected to rsfMRI, at which time they were food-restricted for 21 days. Then rats had to learn a new fixed location of four rewards in the maze for 4 days during the reversal learning phase, during which they again received daily J60 or saline injections. After this, rats were subjected for the final time to rsfMRI, at which time they were food-restricted for 28 days. There were in total five groups of rats. See Table 1 for their differences. B. Food restriction (FR) experiment. Rats were subjected to five rsfMRI scans: i) before FR, ii) after 7 days of FR, iii) after 21 days of FR, iv) after 28 days of FR, and v) 28 days after ending the 28 days of FR.

Cohort 1 consisted of 70 healthy adult male ChAT-Cre (n = 37) and WT (n = 33) Long-Evans rats, which were divided into five groups: three treatment groups expressing hM3Dq DREADDs and mCherry in the medial septum-vertical limb of the diagonal band of Broca (MSDB) and two control groups only expressing mCherry in the MSDB (see Table 1).

**Table 1.** Overview of the five groups in cohort 1. #1: number of animals initially included in the study (excluding the one animal that died after the stereotaxic surgery), #2: number of animals included in the resting-state fMRI analysis.

| Group | hM3Dq DREADD and/or mCherry expression | Specificity of DREADD/mCherry neuronal expression | Administration of JHU37160 (J60, DREADD agonist) or saline (vehicle) | Timing of stimulation for behavioural experiment | Animals (#1/#2) |
| --- | --- | --- | --- | --- | --- |
| cholDREADD/J60/during | DREADD + mCherry | Cholinergic specificity | J60 | During memory training | ChAT-Cre (15/9) |
| cholDREADD/J60/after | DREADD + mCherry | Cholinergic specificity | J60 | After memory training | ChAT-Cre (12/10) |
| DREADD/J60/during | DREADD + mCherry | Nonspecific | J60 | During memory training | ChAT-Cre (5/0)<br>WT (9/0) |
| sham/J60/during | mCherry | Nonspecific | J60 | During memory training | ChAT-Cre (3/1)<br>WT (11/8) |
| sham/saline/during | mCherry | Nonspecific | saline | During memory training | ChAT-Cre (1/1)<br>WT (13/9) |

The two treatment groups with cholinergic specificity consisted entirely of ChAT-Cre rats. The remainder of the ChAT-Cre rats were randomly assigned to the other three groups, where genotype did not matter, as cholinergic specificity was not required in these groups. To achieve DREADD and/or mCherry expression in the MSDB, rats were subjected to an intracranial adeno-associated viral (AAV) injection when approximately four to six months old (mean age at surgery: (140 ± 2) days old, mean weight at surgery: (559 ± 7) g). Minimally six weeks later, allowing for stable DREADD/mCherry expression, rats were subjected to a pharmacological functional connectivity fMRI scanning session while being food-restricted for 7-12 days. Later, animals were subjected to memory training in the radial arm maze, which necessitated food restriction, and which consisted of two learning phases: acquisition learning (10 days) and reversal learning (4 days). On each memory training day, animals received either a J60 or saline injection (see Table 1). In addition, rats were subjected to two resting-state fMRI scans: one after termination of the acquisition learning phase (at 21 days of food restriction) and one after termination of the reversal learning phase (at 28 days of food restriction).

Cohort 2 consisted of 12 healthy adult male ChAT-Cre (n = 9) and WT (n = 3) Long-Evans rats. These animals were not subjected to any kind of surgery. Rats were subjected to five resting-state fMRI scans: i) before food restriction, ii) after 7 days of food restriction (12 g of food per day), iii) after 21 days of food restriction, iv) after 28 days of food restriction, and v) 28 days after ending the food restriction. The animals were approximately four months old at the first resting-state fMRI scanning session (mean age: (121 ± 0) days old, mean weight: (549 ± 10) g).

### Adeno-associated viral (AAV) injection

To achieve DREADD and/or mCherry expression in the MSDB, all animals from cohort 1 received an intracranial AAV injection. Animals were anaesthetised using isoflurane (IsoFlo, Abbott, The Netherlands) in air (induction: 5%, maintenance during drilling: 3%, further maintenance: 2.5%). The head was shaved and then fixed in a stereotactic device (Stoelting Europe, Ireland). Vidisic eye gel (Bausch + Lomb, Belgium) was applied to the eyes, after which they were covered to protect them from the light. The head skin was disinfected using 70% ethanol (MATS bvba, Belgium) and iso-Betadine gel (Mylan, Belgium), after which a topical subcutaneous injection of 0.1 mL 1% lidocaine hydrochloride (Xylocaine 1%, Aspen Pharma Trading Limited, Ireland) was administered. Next, a midline incision was made and the skull exposed. If necessary, the skull was repositioned to a flat skull position with bregma and lambda at the same height. A burr hole was drilled using a handheld electric drill with a 0.9 mm diameter drill tip (Fine Science Tools, Germany) at the following coordinates relative to bregma: ML 1.40 mm (right hemisphere), AP +0.72 mm. A glass capillary micropipette filled with 2.5 μL viral solution was inserted into the brain at an angle of 12° to reach the middle of the MSDB at DV -6.70 mm from the brain surface. A total volume of ca. 2 μL viral solution was injected in 36 steps of 55.2 nL with a one-minute interval and at a speed of 23 nL/s using a Nanoject II microinjector (Drummond Scientific, USA). After removal of the glass capillary micropipette, a subcutaneous injection of 0.025 mg/kg buprenorphine hydrochloride (Temgesic, Indivior, Ireland) was administered and the wound sutured. Finally, 2% lidocaine hydrochloride ointment (Xylocaine 2% gel, Aspen Pharma Trading Limited, Ireland) and iso-Betadine gel were topically applied before placing the animal in a clean home cage on a heating pad or under a heat lamp until recovery from the anaesthesia. One animal did not wake up after the surgery.

Three different viruses were employed in this study. For specific cholinergic neuronal hM3Dq DREADD-mCherry expression, ChAT-Cre rats were injected with pAAV5-hSyn-DIO-hM3D(Gq)-mCherry (Addgene, USA) at a titer of 7.8-8.2×10^12^ genome copies (GC)/mL. For nonspecific neuronal hM3Dq DREADD-mCherry expression, ChAT-Cre or WT rats were injected with pAAV5-hSyn-hM3D(Gq)-mCherry (Addgene) at a titer of 1.5×10^13^ GC/mL. For nonspecific neuronal mCherry expression, ChAT-Cre or WT rats were injected with pAAV5-hSyn-mCherry (Addgene) at a titer of 1.4-2.8×10^13^ GC/mL.

### Memory training in the radial arm maze

Animals from cohort 1 were trained in a custom-made eight-arm radial arm maze to assess visuospatial learning and memory. The radial arm maze consisted of a central regular octagonal zone with each of the eight sides measuring 9.8 cm and eight equidistantly spaced arms, each with a length of 48 cm, an inner width of 8.3 cm and 25.5 cm high walls on all sides. Bird spikes were mounted on the edges to prevent rats from escaping the maze, while ensuring optimal vision for the rats of the surrounding environment. Both two-dimensional and three-dimensional visual cues surrounded the maze. The maze was placed behind a wall so that the researcher was not visible to the rat while in the maze. Tests were performed in a dimly lit room with an additional infrared illuminator (Stoelting Europe, Ireland). The maze had been made of black material, but for the purposes of this study, the floor was made white in order to achieve optimal tracking of the Long-Evans rats. Live tracking of the centre of the animal (the Long-Evans rats’ black hood) was done with the ANY-maze video tracking software (ANY-maze, Stoelting Europe, Ireland) through an overhead infrared USB camera (Stoelting Europe, Ireland). Rewards consisted of half honey pops. At all times, rewards were hidden in cups underneath the end of all arms and inaccessible to the animals by means of a metal mesh cover, to provide potential olfactory stimulation to the animals in all arms. Depending on the testing phase, some or all the arms contained rewards that were accessible to the rats. These rewards were placed on the metal mesh, but still below floor level, to rule out visibility from the central zone of the maze.

The test consisted of four phases: a familiarisation phase, a habituation phase, an acquisition phase and a reversal phase. The last two phases (the learning phases) always took place during the afternoon. In the familiarisation phase, animals were familiarised with the reward in their home cage and with the training personnel (animal handling) for five consecutive days. Next, during the habituation phase, animals were habituated to the maze for five to seven consecutive days. During this phase, rewards were accessible in all arms. During the first two days, rewards were put visibly at the end of the arms. After this, rewards were hidden at the end of the arms below floor level. Each day consisted of a single trial of max. 15 min or until all rewards had been found and eaten. Between trials the maze was thoroughly cleaned with 70% propanol (IPASEPT 70, VWR Chemicals, Belgium) to remove odour traces from the previous animal. If animals were able to find a majority of the rewards (five or more out of eight) by the end of the fifth day, the habituation phase came to an end. If not, the habituation phase continued for two more days. In the rare case that animals were still not able to locate the rewards by the end of the seventh day, animals were excluded from the memory training experiment (n = 4). During the acquisition phase, which lasted ten days, rewards were accessible to the rats in four arms (the same arms for all animals). Finally, during the reversal phase, which lasted four days, the position of the accessible rewards was changed to the other four arms of the maze. Each day of the acquisition and reversal phases consisted of three trials per animal, each lasting max. 5 min (or until all rewards had been found and eaten). During both the acquisition and reversal phases, all groups except the one group that received MSDB stimulation after the training (cholDREADD/J60/after), received a subcutaneous injection of 0.1 mg/kg JHU37160 dihydrochloride (J60, Hello Bio Ltd, UK) or saline on each training day 15 min before the first trial started. The second and third trial took place 45 min and 75 min post-injection, respectively. During each trial, animals were placed in the centre of the maze, but each time with a random starting orientation (nose pointing in a different direction) in order to prevent egocentric learning and promote allocentric learning. The group that received specific cholinergic MSDB stimulation after training received a subcutaneous injection of 0.1 mg/kg J60 approximately 15 min before the light phase started at the end of the day (on average (22 ± 1) min after finishing their last trial). From the first day of the habituation phase onwards, rats were food-restricted to motivate them to look for the rewards. On days with trials in the maze rats received 10 g of standard food pellets (on top of the rewards that were eaten), while on days when no trials took place rats received 12 g of food to compensate for the lack of food rewards. We measured the *ad libitum* daily food consumption in six healthy adult (five-to-six-month-old) male Long-Evans rats for four days and observed that they consumed on average (31 ± 1) g of food per day.

The following parameters were measured using ANY-maze software: time spent and distance travelled in the entire maze and per zone (one centre zone + eight arm zones), number of entries in each zone (defined as the crossing of the animal’s centre into the zone), latency to first entry into each zone and average speed in the maze. In addition, we calculated the % reference memory errors (defined as the number of entries in non-baited arms divided by the total number of arm entries x100), the % distance travelled in non-baited arms (sum of the distance travelled in each of the four non-baited arms divided by the total distance travelled in the maze x100), the % time spent in non-baited arms (sum of the time spent in each of the four non-baited arms divided by the total time spent in the maze x100), latency to first entry in a non-baited arm (shortest latency to entry in any one of the four non-baited arms), latency to first entry in a baited arm (shortest latency to entry in any one of the four baited arms), and the % working memory errors (number of arm re-entries divided by the total number of arm entries x100). For each acquisition or reversal training day, an average of the three trials per day was calculated per parameter and used for further analysis. The following parameters were subjected to statistical analysis: average total duration of the three trials, average total distance travelled during the three trials, mean speed in the maze, % reference memory errors, % time spent in non-baited arms, % distance travelled in non-baited arms, latency to entry in first non-baited arm, latency to entry in first baited arm and % working memory errors.

Cohort 1 was subdivided into seven batches. Due to COVID-19-related reasons, the last batch could not be included in the memory training analysis. Some animals had to be excluded from the memory training since they did not learn to locate or had no interest in rewards during the habituation phase (n = 4). Other animals were excluded based on outlier analysis (n = 3). One animal of the DREADD/J60/during group died on the second day of acquisition training following injection of J60 and was therefore excluded from the analysis. In the end, a total of 48 animals were included in the memory training analysis: 9 cholDREADD/J60/during rats, 10 cholDREADD/J60/after rats, 10 DREADD/J60/during rats, 9 sham/J60/during rats and 10 sham/saline/during rats.

### Pharmacological functional connectivity fMRI – cohort 1

To assess the effects of MSDB activation on resting-state networks, rats from cohort 1 were subjected to pharmacological functional connectivity fMRI with the DREADD agonist J60 or its vehicle saline. Rats were food-restricted (receiving 12 g of standard food pellets per day) for 7-12 days prior to scanning in order to have similar conditions as during memory training. Functional MRI data for functional connectivity analysis were acquired on a 7T PharmaScan MRI system (Bruker, Germany) with Paravision 6 software with a similar set-up, anaesthesia protocol, scanning sequence, orientation and dimensions as described before for resting-state fMRI^45^. Magnetic field inhomogeneity was corrected by local shimming in an ellipsoid volume of interest within the brain. Functional MRI was acquired using a single shot gradient echo EPI-sequence (TR 2000 ms, TE 20 ms, FOV (30 x 30) mm^2^, matrix [128 x 128], 18 coronal slices of 0.7 mm + 0.1 mm slice gap (covering the cerebrum and excluding the olfactory bulb)). Rats were anesthetized with isoflurane in a mixture of O2 (30%) and N2 (70%) (5% induction; Forene; Abbott, Belgium) after which a subcutaneous (s.c.) bolus injection of 0.05 mg/kg medetomidine hydrochloride (Domitor, Pfizer, Karlsruhe, Germany) was administered to sedate the animals. Fifteen minutes post-bolus-injection a continuous s.c. infusion of 0.1 mg/kg/h medetomidine was started. Following bolus injection, isoflurane was gradually decreased to 0.4% during the fMRI scan. At the end of the scan session, a subcutaneous injection of 0.1 mg/kg atipamezole (Antisedan®, Pfizer, Germany) was administered to counteract the effects of the medetomidine anesthesia.

Functional MRI scanning started at 35 min post-medetomidine bolus. After 10 min of baseline resting-state fMRI acquisition, 0.1 mg/kg J60 or saline (1 mL/kg) was infused in the rat via a previously prepared subcutaneous infusion line over a timespan of one minute and an additional 60 min of fMRI data were acquired. Hence, the total duration of this pharmacological functional connectivity fMRI scan was 70 min and a total of 2100 volumes were acquired.

Image preprocessing was performed in Advanced Normalization Tools (ANTs) and SPM12 in MATLAB 2014a (MathWorks, USA). A study-specific echo planar imaging (EPI) template was made in ANTs based on the first volume of all 69 subjects that received a pharmacological functional connectivity fMRI scan. Pharmacological functional connectivity fMRI images within each session were realigned to the first image using a combined affine and diffeomorphic normalisation in ANTs. The remainder of the analysis was performed in SPM12. All EPI datasets were normalised to the study-specific EPI template using an affine transformation followed by the estimation of the nonlinear deformations. Next, in-plane smoothing was performed using a Gaussian kernel with full width at half maximum (FWHM) of twice the voxel size (FWHM (0.468 × 0.468 × 0.8) mm^3^). Finally, datasets were filtered using the Resting State fMRI Data Analysis toolbox (REST1.8). The band-pass filter was set between 0.01 and 0.1 Hz to retain the low frequency fluctuations of the blood-oxygen level dependent (BOLD) signal time course.

For each scan, the data were split into seven sets of 300 volumes (10 min). The first 10 min constituted the baseline resting-state fMRI data, followed by six post-J60/saline 10 min time intervals. The first 10 min after subcutaneous infusion of J60/saline were disregarded.

Region of interest (ROI)- and seed-based functional connectivity (FC) analyses were performed as described before for resting-state fMRI data^45^. In this study, we investigated the following ROIs (left and right taken together as a single ROI) and seeds (the left hemisphere ROIs): orbitofrontal cortex (OF), prelimbic-infralimbic cortex (PrL-IL) and anterior cingulate cortex (Ant Cg) (main constituents of the anterior default mode-like network (DMLN)); posterior cingulate cortex (Post Cg), retrosplenial cortex (RS), parietal association cortex (PtA) and posterior parietal cortex (PtP) (main constituents of the posterior DMLN); anterior hippocampus proper (Ant Hc), posterior hippocampus proper (Post Hc) and dentate gyrus (DG) (main constituents of the hippocampus); primary motor cortex (M1) and primary somatosensory cortex (S1) (main constituents of the lateral cortical network (LCN), analogous to the task-positive network (TPN) in humans); nucleus accumbens (Acb), ventral caudate putamen (CPu) and insular cortex (Ins) (main constituents of the salience-like network (SN)). The anterior DMLN, posterior DMLN, LCN and SN were the four most prominent bilateral resting-state network modules revealed by independent component analysis of the resting-state fMRI data (10 min baseline) (data not shown). In addition to these four network modules, we investigated the hippocampus, which is especially relevant for spatial memory. These ROIs/seeds were defined as square regions of four voxels using MRIcron software, based on the Paxinos and Watson rat brain atlas (6^th^ edition), and delineated on the study-specific EPI template.

#### ROI-based FC analysis

The filtered functional connectivity fMRI datasets and a mask for each ROI were used as an input to REST to extract the time courses for each ROI from each subject. Correlation coefficients between the time courses of each pair of ROIs were calculated and z-transformed using an in-house MATLAB program. The average group z-transformed correlation values were presented in a FC strength matrix. The FC strength within and between network modules was computed by calculating the average of the pairwise correlation values between all pairs of ROIs which the network modules consisted of (as listed above).

#### Seed-based FC analysis

Seed-based FC analyses were performed for each of the above-mentioned seeds (left hemisphere ROIs) except M1 and S1 since ROI-based FC analysis showed no significant differences within the LCN (M1-S1 connection). BOLD signal time courses for each of these seeds were extracted from all filtered datasets using the REST toolbox. These were used in SPM12 in a generalised linear model to compare them to the time courses of all other voxels in the brain. Individual statistical FC maps for all animals were obtained, which represent the voxels in the brain that significantly correlate with the temporal signal of the seed region. Mean FC maps for each seed region were computed per group and per 10 min time interval (one-sample t-test, uncorrected, p≤0.001, minimal cluster size = 10 voxels). These contain all clusters of 10 voxels or more that are significantly correlated with the seed region at group-level. The group-averaged FC maps were saved as masks and a summed mask (summed across all groups and time intervals) was created. For each subject at each timepoint the mean T-value (measure of FC strength) of the individual FC map was extracted within this summed mask.

Two animals were identified as outliers in JMP Pro (>1.5 times the interquartile range when plotting the data as box plots) and therefore excluded from these analyses. A total of 67 animals were included in the pharmacological functional connectivity fMRI analysis: 27 cholDREADD/J60 rats, 14 DREADD/J60 rats, 14 sham/J60 rats and 12 sham/saline rats.

### Resting-state fMRI – cohort 1 and 2

Resting-state fMRI was acquired exactly as described above for the first 10 min of the pharmacological functional connectivity fMRI. 300 volumes (10 min) were acquired per resting-state fMRI session. No J60 or saline was administered. Image preprocessing was performed completely in SPM12 in MATLAB. Resting-state fMRI images within each session were realigned to the first image using a least-squares approach and a 6-parameter (rigid body) spatial transformation. All other preprocessing steps were performed exactly as described above. The same EPI template was used for the normalisation of the data. ROI- and seed-based FC analyses were performed as described above with the same set of ROIs/seeds.

#### Memory experiment (cohort 1)

To assess the effects of memory training in the radial arm maze (both acquisition and reversal training) with or without MSDB stimulation on resting-state networks, rats from cohort 1 were subjected to resting-state fMRI scans i) before learning (the first 10 min of the pharmacological fMRI scan described above), ii) after acquisition training and iii) after reversal training. All three resting-state fMRI data sets were acquired and processed in the same way. The first 10 min of the pharmacological functional connectivity fMRI scan from above served not only as baseline for the acquired post-J60 functional connectivity fMRI data, but also as a pre-learning baseline for the other two post-learning resting-state fMRI data sets. As mentioned above, at this first timepoint rats had been food-restricted (12 g of food per day) for 7-12 days. The second resting-state fMRI scan was acquired after animals finished the acquisition phase of the memory training in the radial arm maze. At this post-acquisition timepoint, animals had been food-restricted (10 g of food on days with habituation/memory training trials in the maze, 12 g of food on days without trials in the maze) for 21 days. The third and final resting-state fMRI scan was acquired after animals finished the reversal phase of the memory training in the radial arm maze. At this post-reversal timepoint, animals had been food-restricted (10 g of food on days with trials in the maze, 12 g of food on days without trials in the maze) for 28 days. Only the scans from the animals that were included in the memory training analysis were included in the resting-state fMRI analysis. Moreover, 70% of the rats (7/10) that received nonspecific MSDB stimulation via DREADDs did not finish the acquisition phase of the memory training (see later) and were therefore not subjected to post-acquisition or post-reversal resting-state fMRI scans. In the end, only three animals from this group received post-learning fMRI scans. This number was deemed insufficient for statistical analysis. This particular group was therefore excluded from the resting-state fMRI analysis. In the end, a total of 38 animals were included in the resting-state fMRI analysis: 9 cholDREADD/J60/during rats, 10 cholDREADD/J60/after rats, 9 sham/J60/during rats and 10 sham/saline/during rats (see Table 1).

#### Food restriction experiment (cohort 2)

To assess the effects of chronic food restriction on resting-state networks, rats from cohort 2 were subjected to resting-state fMRI scans i) before food restriction, ii) after 7 days of food restriction (12 g of food per day), iii) after 21 days of food restriction, iv) after 28 days of food restriction, and v) 28 days after ending the food restriction. Twelve rats were included in this experiment.

### Statistical analysis

All data sets were subjected to linear mixed model analysis. Both random intercept and random slope models were fitted, and a likelihood ratio test was performed to compare the goodness of fit of the two statistical models. If addition of a random slope significantly improved the model, then the random slope model was used. If not, the simpler random intercept model was chosen to analyse the data.

For the memory training analysis, the acquisition phase data and the reversal phase data were analysed separately. In these analyses, random slope models were used, and time was considered a continuous variable. In case of a significant main group effect, post-hoc analysis was performed using a Dunnett’s multiple comparisons with control test with the “sham/J60/during” group as control group. In case of a significant group*time interaction, linear mixed models were made split by group to investigate the effect of time per group. The resulting time effects of the different groups were false discovery rate (FDR) corrected.

For the fMRI analyses, random intercept models were used, and time was considered a categorical variable as to be able to compare post-J60 or post-learning timepoints with the baseline timepoint. The resulting main group effects, time effects and group*time interactions of all the investigated functional connections were false discovery rate (FDR) corrected. If the FDR-corrected group effect for a given connection was significant and there was no significant FDR-corrected group*time interaction, then post-hoc analysis was performed using a Dunnett’s multiple comparisons with control test with the “sham/J60/during” group as control group. If the FDR-corrected time effect for a given connection was significant and there was no significant FDR-corrected group*time interaction, then post-hoc analysis was performed using a Dunnett’s multiple comparisons with control test with the baseline timepoint as control timepoint. In the case of a significant FDR-corrected group*time interaction, linear mixed models were made split by group (to investigate the effect of time per group) and split by time (to investigate the effect of group per timepoint). Respectively, the resulting time effects of the different groups and the group effects of the different timepoints were FDR corrected. If still significant, then a Dunnett’s multiple comparisons with control test was run with respectively the baseline timepoint as control timepoint or with the “sham/J60/during” group as control group.

For the analysis of the resting-state fMRI data from the food restriction experiment (cohort 2), random intercept models were fitted with only time as a main effect. These were FDR-corrected and if significant, post-hoc analysis of the main time effect was performed in two ways: i) comparing all timepoints with the pre-food restriction baseline and ii) comparing all timepoints with the 7-day food restriction timepoint. All statistical analyses were performed in JMP Pro 14. Statistical significance was set at p≤0.05. The graphs were made in GraphPad Prism 8. Data are represented as mean ± standard error of the mean.

## Results

### Memory training in the radial arm maze

All investigated parameters showed a significant effect of time (mean speed during acquisition training: p=0.0051, latency to entry in first baited arm during reversal training: p=0.0142, all other investigated parameters during acquisition and reversal training: p < 0.0001). This indicates that all rats that participated in this task showed a significant improvement over time, i.e. they all learned the task, both the initial acquisition and the reversal (Fig.2).

**Figure 2.**
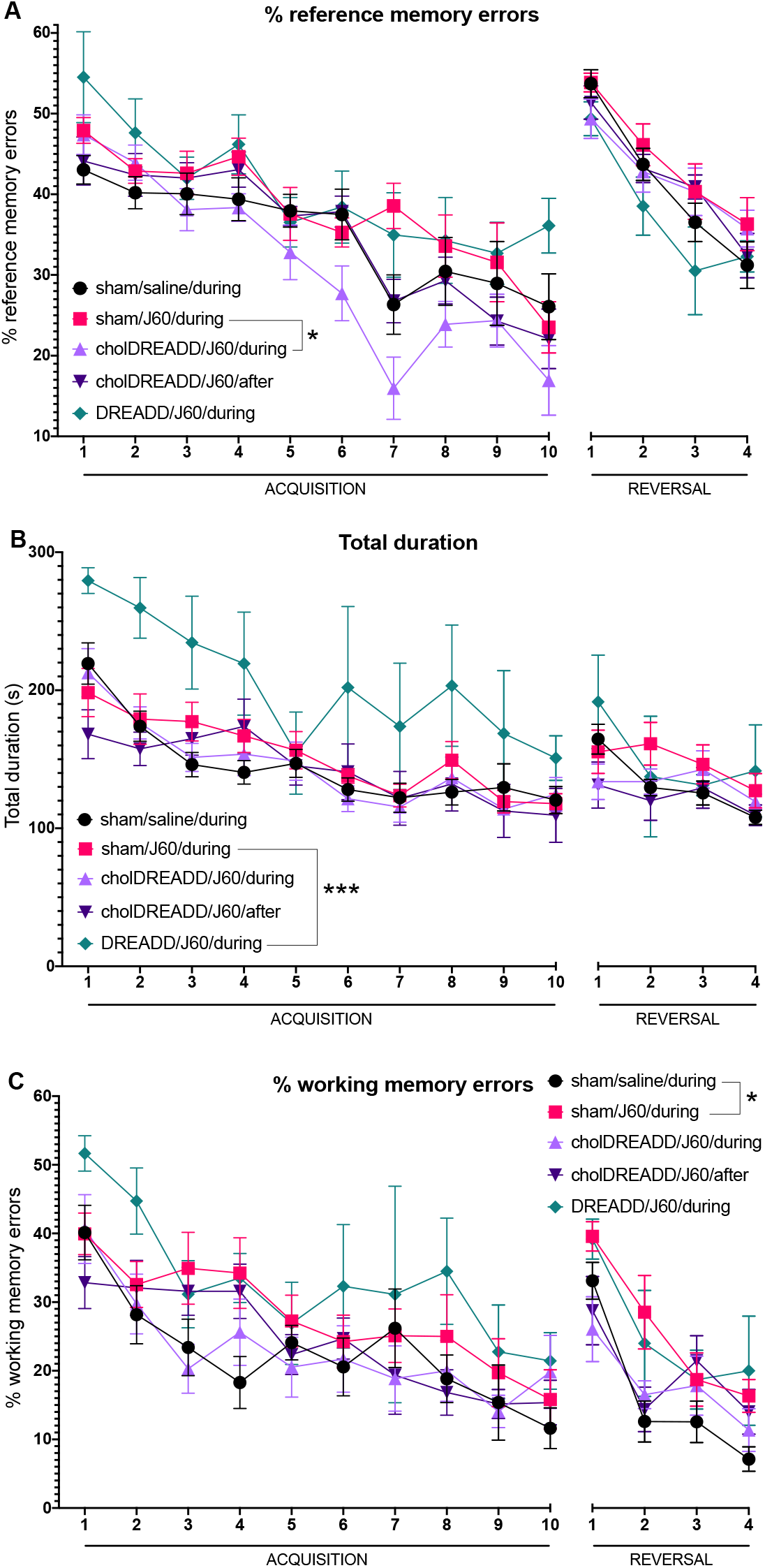
Memory training in the radial arm maze: acquisition and reversal learning. A. Rats receiving specific cholinergic medial septum-diagonal band of Broca (MSDB) stimulation during memory training (cholDREADD/J60/during, light purple ▲) had a significantly lower % reference memory errors than J60-treated sham rats (sham/J60/during, fuchsia ▄) during acquisition, but not during reversal. B. Rats receiving nonspecific neuronal MSDB stimulation during memory training (DREADD/J60/during, pine green ◆) took significantly more time to finish the acquisition trials than J60-treated sham rats (sham/J60/during, fuchsia ▄). C. J60-treated sham rats (sham/J60/during, fuchsia ▄) had a significantly higher % working memory errors than saline-treated sham rats (sham/saline/during, black ●) during reversal, but not during acquisition training. Sham/saline/during: n = 10; sham/J60/during: n = 9; cholDREADD/J60/during: n = 9; cholDREADD/J60/after: n = 10; DREADD/J60/during: acquisition day 1: n = 10, day 2: n = 8, day 3: n = 5, day 4: n = 4, acquisition day 5 till end of reversal training: n = 3. *p≤0.05, ***p≤0.001

During acquisition training, significant group effects were observed for all the investigated parameters (see below) except the average total distance travelled in the maze (p=0.2744) and the % time spent in non-baited arms (p=0.0798). No significant group*time interactions were observed (p>0.05), indicating that all groups follow the same temporal changes. Post-hoc analysis showed that the differences were either between cholDREADD/J60/during and sham/J60/during rats or between DREADD/J60/during and sham/J60/during rats (see below).

During reversal training, significant group differences were observed in only two of the investigated parameters: % time spent in non-baited arms and % working memory errors (number of arm re-entries divided by the total number of arm entries x100) (see below). Only one parameter showed a significant group*time interaction, i.e. the average total distance travelled in the maze (p=0.0216). Post-hoc analysis showed that significant FDR-corrected time effects were observed in all groups except the group that received nonspecific neuronal MSDB stimulation during training (sham/saline/during: p=0.0003, sham/J60/during: p=0.0001, cholDREADD/J60/during: p=0.0111, cholDREADD/J60/after: p=0.0013, DREADD/J60/during: p=0.0804).

The complete statistical analysis can be found in Supplementary Tables 1, 2 and 3.

By the end of the acquisition training (day 10), rats had an average weight of (86.3 ± 0.4) % of their baseline weight on the first day of the food restriction. By the end of the reversal training (day 4), rats had reached an average weight of (83.6 ± 0.5) % of their baseline weight.

### Specific cholinergic neuronal MSDB stimulation during training improved spatial reference memory during acquisition in the radial arm maze

Rats that received specific cholinergic neuronal MSDB stimulation during training (i.e. a subcutaneous injection of J60 15 min before the first trial) showed a significantly lower % reference memory errors (group effect: p=0.0068; post-hoc adjusted p-value for the difference with sham/J60/during rats = 0.0150; Fig.2A), a significantly lower % distance travelled in non-baited arms (p=0.0103: adj. p=0.0129) and a significantly longer latency to entry in first non-baited arm (p=0.0346; adj. p=0.0199) than J60-treated sham rats during acquisition training. All these measures pertain to reference memory, i.e. remembering which arms are baited or not. Hence, a significant improvement of reference memory was observed after specific cholinergic neuronal MSDB stimulation during acquisition training. This positive effect on reference memory specific for cholinergic neuronal MSDB stimulation during memory training, however, was not observed during reversal training. The timing of the stimulation also proved to be important, since a similar positive effect on reference memory was not observed in rats that received specific cholinergic neuronal MSDB stimulation after training.

### Nonspecific neuronal MSDB stimulation during training resulted in aberrant behavior

Most of the rats that received nonspecific neuronal MSDB stimulation during training displayed abnormal behaviour. Many rats in this group seemed to exhibit a pronounced loss of appetite following J60 injection, disregarding food rewards as well as daily food pellets, despite being food-restricted. These rats exhibited a significantly lower speed in the maze and several rats also showed a pronounced increase in water consumption. Rats that did not eat all their food during the 24 h between daily training sessions were immediately excluded from the behavioural and resting-state fMRI study (n = 7/10). While on day 1 of the acquisition training 10 rats in this group participated in the behavioural experiment, on day 2 only 8 rats still participated, on day 3 only 5 rats, on day 4 only 4 rats and from day 5 onwards only 3 rats. Hence, there was a 70% dropout rate in this group by day 5 of the acquisition training. There were no dropouts in any of the other groups. The statistical analysis was performed on all acquired data and also the graphs include all acquired data until exclusion from the study.

Rats that received nonspecific neuronal MSDB stimulation during training had a significantly higher average total duration of trials in the maze (group effect: p<0.0001; post-hoc adjusted p-value for the difference with sham/J60/during rats = 0.0003; Fig.2B), a significantly lower mean speed in the maze (p=0.0024: adj. p=0.0011) and a significantly longer latency to entry in first baited arm (p=0.0074: adj. p=0.0106) compared with J60-treated sham rats during acquisition training. During reversal training, these rats spent a higher % time in non-baited arms than the J60-treated sham rats (p=0.0148: adj. p=0.0142).

Hence, we can conclude that nonspecific neuronal MSDB stimulation led to undesired side effects and had no positive effect on learning and memory as far as this could be evaluated.

### J60-treated sham rats had a worse spatial working memory during reversal in the radial arm maze than saline-treated sham rats

Saline-treated sham rats had a significantly lower % working memory errors than J60-treated sham rats during reversal training (group effect: p=0.0229; post-hoc adjusted p-value for the difference with sham/J60/during rats=0.0129, Fig.2C). While there was also a significant group effect for the % working memory errors during acquisition training (p=0.0247), post-hoc analysis revealed no difference between J60-treated sham rats and any of the other groups.

#### Pharmacological functional connectivity fMRI reveals a differential impact of acute specific cholinergic and nonspecific neuronal MSDB stimulation on resting-state networks

### Region of interest-based functional connectivity analysis revealed decreased default mode-like network connectivity after both acute specific cholinergic and nonspecific neuronal MSDB stimulation

Functional connectivity (FC) within five network modules were investigated, namely the anterior default mode-like network (DMLN), posterior DMLN, hippocampus (see next section), lateral cortical network (LCN), and salience-like network (SN). The complete statistical analysis can be found in Supplementary Tables 4, 5 and 6.

No significant differences were observed within the LCN (p>0.05) (Supplementary Table 4). A significant FDR-corrected group effect was observed for the FC within the SN (p=0.0095), but post-hoc analysis revealed no significant differences between J60-treated shams and any of the other groups (p>0.05) (Supplementary Table 4).

For the FC within the anterior DMLN and posterior DMLN a significant FDR-corrected group*time interaction was observed (respectively p=0.0062 and p<0.0001) (Supplementary Table 4).

FC within the anterior DMLN changed significantly over time in rats that received specific cholinergic neuronal MSDB stimulation (FDR-corrected p=0.0020) and in rats that received nonspecific neuronal MSDB stimulation (FDR-corrected p=0.0156), but not in the sham groups (both p>0.05) (Fig.3 and 4, Supplementary Table 5). The rats that received specific cholinergic MSDB stimulation showed a significantly lower FC at 10-20 min (post-hoc adjusted p-value for the difference with sham/J60 rats = 0.0026), 20-30 min (adj. p=0.0041) and 30-40 min post-J60-injection (p.i.) (adj. p=0.0206) compared with baseline (10 min pre-J60), but not during the later time intervals (40-50 min and 50-60 min p.i.: p>0.05). The rats that received nonspecific MSDB stimulation had a significantly lower FC at 10-20 min p.i. (adj. p=0.0375) compared with baseline, but not during later time intervals (p>0.05). The post-hoc analysis per time interval showed significant FDR-corrected group effects for the 10-20 min and 20-30 min p.i. time intervals (respectively p=0.0024 and p=0.0114), but not during any other time interval (baseline, 30-40 min, 40-50 min and 50-60 min pi.: p>0.05) (Supplementary Table 6). The specifically stimulated rats (cholDREADD/J60) had a significantly lower FC than J60-treated sham rats during the 10-20 min and 20-30 min p.i. time intervals (respectively adj. p=0.0006 and 0.0016). The nonspecifically stimulated rats (DREADD/J60) had a significantly lower FC than J60-treated sham rats during the 10-20 min p.i. time interval (adj. p=0.0009). Saline-treated sham rats did not differ from J60-treated sham rats (p>0.05).

**Figure 3.**
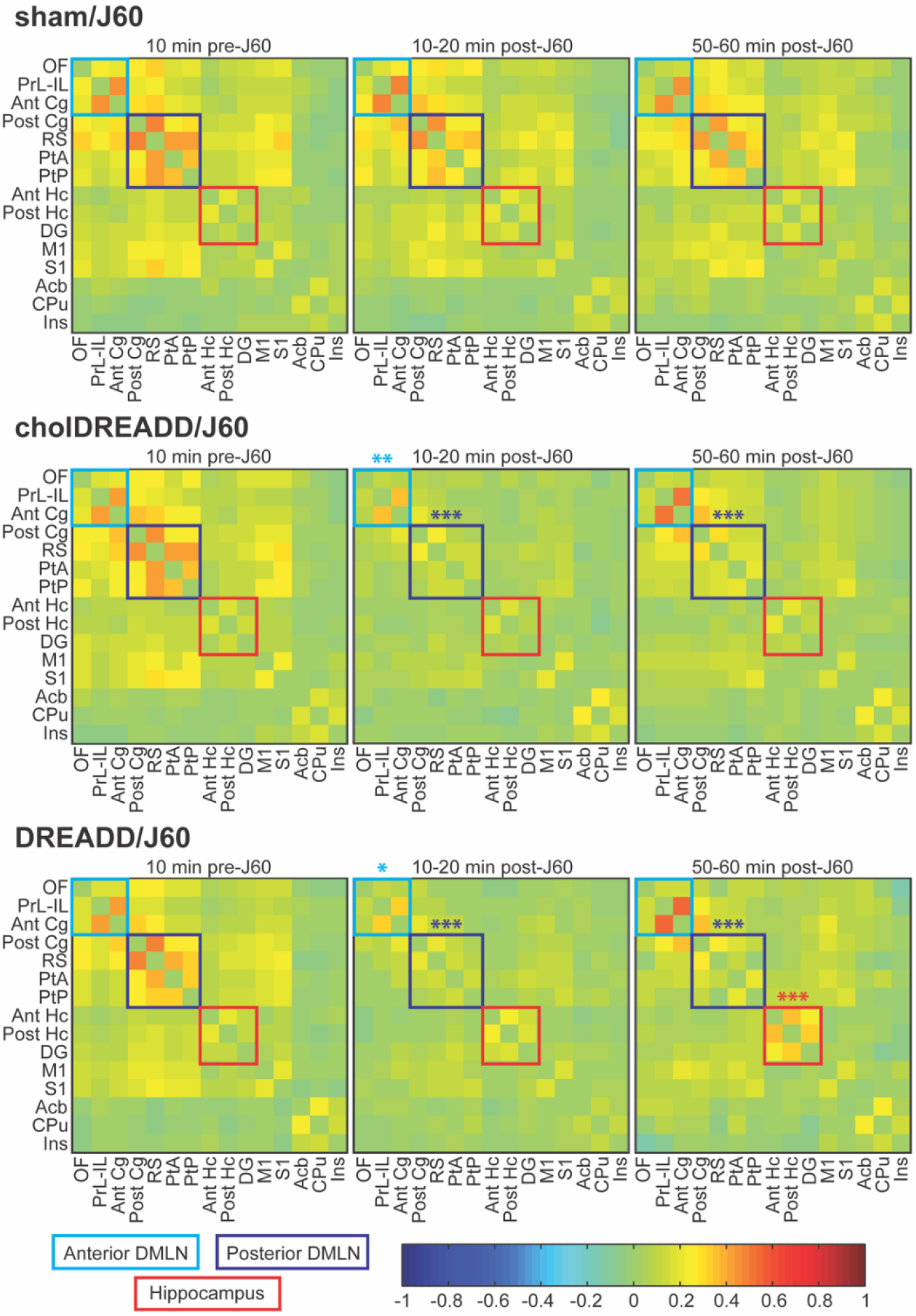
Region of interest (ROI)-based functional connectivity (FC) analysis of the pharmacological fMRI data with the DREADD agonist J60 as acute pharmacological challenge. Group-averaged z-transformed FC matrices with the following ROIs from top to bottom (and from left to right): orbitofrontal cortex (OF), prelimbic and infralimbic cortices (PrL-IL), anterior cingulate cortex (Ant Cg), posterior cingulate cortex (Post Cg), retrosplenial cortex (RS), parietal association cortex (PtA), posterior parietal cortex (PtP), anterior hippocampus proper (Ant Hc), posterior hippocampus proper (Post Hc), dentate gyrus (DG), primary motor cortex (M1), primary somatosensory cortex (S1), nucleus accumbens (Acb), ventral caudate putamen (CPu) and insular cortex (Ins). Cortical ROIs that constitute the anterior default mode-like network (DMLN) and posterior DMLN are delineated by a light blue box and dark blue box, respectively. Hippocampal ROIs are delineated by a red box. Three timepoints are shown here for three of the groups: 10 min before subcutaneous J60 injection (baseline), 10-20 min after J60 injection and 50-60 min after J60 injection. There were no significant changes in the resting-state networks (RSNs) in J60-treated sham rats (sham/J60, top row) and saline-treated sham rats (sham/saline, not shown) over time. Both specific cholinergic (cholDREADD/J60) and nonspecific neuronal medial septum-diagonal band of Broca (MSDB) stimulation (DREADD/J60) led to a pronounced acute decrease in FC within the DMLN (light blue box), which recovered partly over time (middle and bottom rows). Only nonspecific MSDB stimulation (DREADD/J60) led to increased intrahippocampal FC (red box) over time (bottom row). The colour scale indicates the z-transformed correlation values. Sham/J60: n = 14, cholDREADD/J60: n = 27, DREADD/J60: n = 14. *p≤0.05, **p≤0.01, ***p≤0.001. Only the significant changes within a group versus pre-J60 baseline are indicated on these graphs.

FC within the posterior DMLN also changed significantly over time in cholDREADD/J60 and DREADD/J60 rats (both FDR-corrected p<0.0001) and not in the sham groups (both p>0.05) (Fig.3 and 4, Supplementary Table 5). Both the specifically stimulated and nonspecifically stimulated rats had a significantly lower FC at all post-J60 time intervals compared with baseline (adj. p<0.0001 for all). The post-hoc analysis per time interval showed significant FDR-corrected group effects for all post-J60 time intervals (adj. p<0.0001 for all except the last time interval, 50-60 min p.i.: adj. p=0.0006) and not for baseline (p>0.05) (Supplementary Table 6). At all post-J60 time intervals, the specifically stimulated and nonspecifically stimulated rats had a significantly lower FC than J60-treated sham rats (10-20 min p.i.: cholDREADD adj. p<0.0001, DREADD adj. p=0.0001; 20-30 min p.i.: cholDREADD adj. p<0.0001, DREADD adj. p=0.0002; 30-40 min p.i.: cholDREADD adj. p=0.0004, DREADD adj. p=0.0004; 40-50 min p.i.: cholDREADD adj. p=0.0039, DREADD adj. p=0.0013; 50-60 min p.i.: cholDREADD adj. p=0.0154, DREADD adj. p=0.0039). Saline-treated sham rats did not differ from J60-treated sham rats (p>0.05).

**Figure 4.**
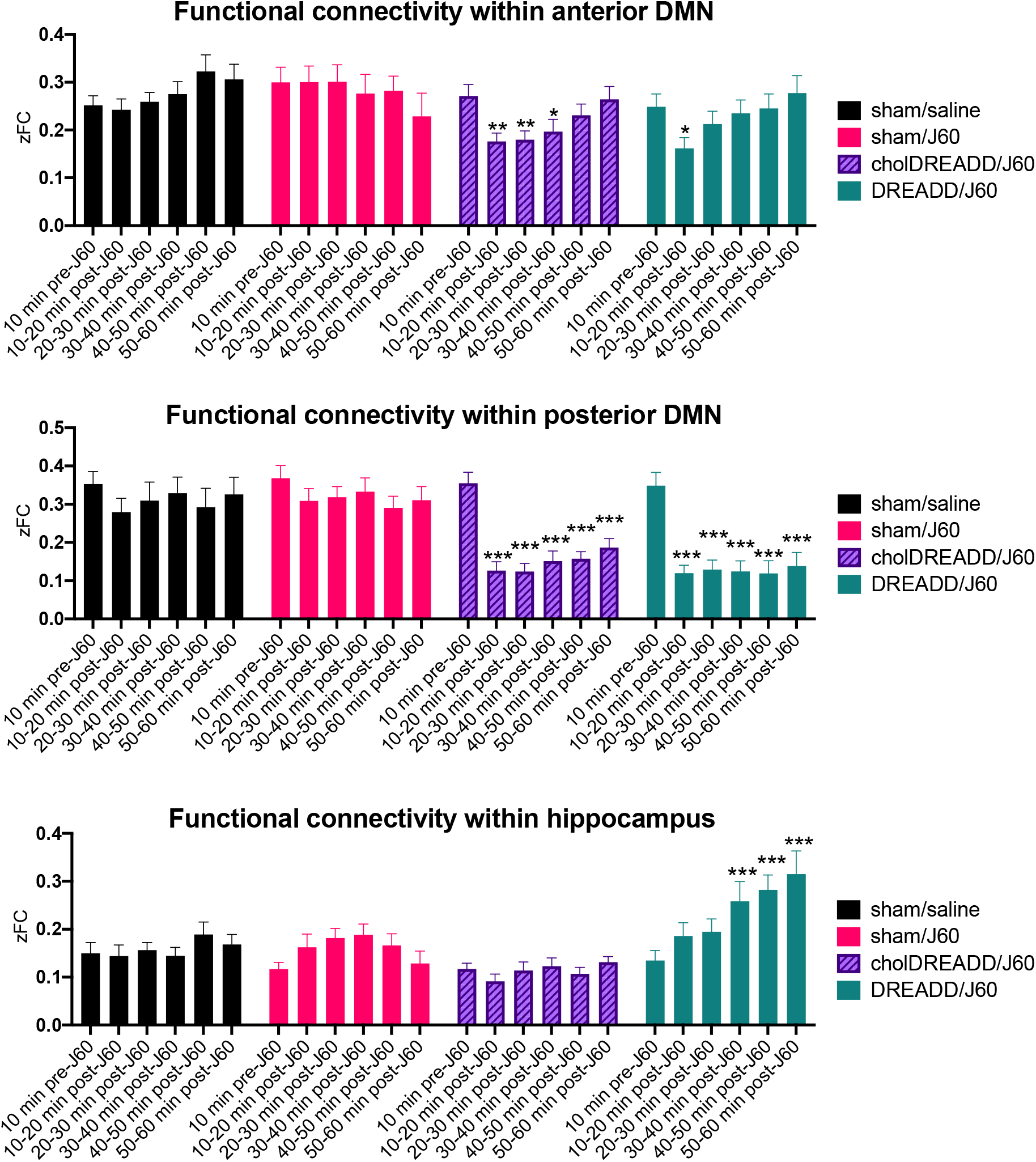
Region of interest (ROI)-based functional connectivity (FC) analysis of the pharmacological fMRI data with the DREADD agonist J60 (or saline) as acute pharmacological challenge. Statistical analysis of the group-averaged z-transformed FC matrices from figure 3. Five network modules were investigated: anterior default mode-like network (DMLN), posterior DMLN, hippocampal network, lateral cortical network (LCN) and salience-like network (SN). No significant changes were shown in the latter two networks (data not shown). Top panel: Both specific cholinergic (cholDREADD/J60, purple) and nonspecific neuronal medial septum-diagonal band of Broca (MSDB) stimulation (DREADD/J60, pine green) resulted in an acute decrease in FC in the anterior DMLN, which recovered over time. Middle panel: Both specific cholinergic (cholDREADD/J60, purple) and nonspecific neuronal MSDB stimulation (DREADD/J60, pine green) resulted in an acute and persistent decrease in FC in the posterior DMLN. Bottom panel: Only nonspecific neuronal MSDB stimulation (DREADD/J60, pine green) resulted in increased intrahippocampal FC over time. Sham/saline: n = 12, sham/J60: n = 14, cholDREADD/J60: n = 27, DREADD/J60: n = 14. *p≤0.05, **p≤0.01, ***p≤0.001. Only the significant changes within a group versus pre-J60 baseline are indicated on these graphs.

While FC in the posterior DMLN showed a very pronounced acute decrease after MSDB stimulation (both specific and nonspecific) and remained decreased for the entire scanning session, FC in the anterior DMLN showed a less pronounced acute decrease after MSDB stimulation and recovered completely within one hour after J60 injection (Fig.3 and 4).

### ROI-based FC analysis revealed increased intrahippocampal connectivity after acute nonspecific neuronal MSDB stimulation

For the FC within the hippocampus a significant FDR-corrected group*time interaction was observed (p<0.0001) (Supplementary Table 4).

Post-hoc analysis per group revealed a significant change over time in intrahippocampal FC in the rats that received nonspecific neuronal MSDB stimulation (FDR-corrected p<0.0001) but not in any other group (p>0.05) (Fig.3 and 4, Supplementary Table 5). Specifically, the rats receiving nonspecific MSDB stimulation had an increased FC at 30-40 min p.i. (p=0.0005), 40-50 min and 50-60 min p.i. (both p<0.0001) compared with baseline.

Post-hoc analysis per time interval revealed a significant difference in intrahippocampal FC between the groups for every post-J60 time interval and not during baseline (baseline: FDR-corrected p=0.4810, 10-20 min p.i.: p=0.0147, 20-30 min p.i.: p=0.0245, 30-40 min p.i.: p=0.0028, 40-50 and 50-60 min p.i.: p<0.0001) (Supplementary Table 6). Specifically, during the first two post-J60 time intervals animals that received specific cholinergic MSDB stimulation had a lower intrahippocampal FC than J60-treated sham rats (10-20 min p.i.: adjusted p=0.0492 and 20-30 min p.i.: adj. p=0.0494). During the last two post-J60 time intervals nonspecifically stimulated rats showed an increased intrahippocampal FC compared with J60-treated sham rats (40-50 min p.i.: adj. p=0.0031 and 50-60 min p.i.: adj. p<0.0001). At 30-40 min p.i., there was no significant difference between J60-treated sham rats and any other group (p>0.05).

### Seed-based FC analysis confirmed ROI-based FC analysis and revealed increased connectivity with nucleus accumbens after acute specific cholinergic neuronal MSDB stimulation

The seed-based FC analysis largely confirmed the ROI-based FC analysis (see Supplementary Tables 7, 8 and 9 for all p-values). Animals that received MSDB stimulation (both specific and nonspecific) showed a decreased FC after J60 injection with seeds from the anterior DMLN and posterior DMLN: orbitofrontal cortex, anterior cingulate cortex, posterior cingulate cortex, retrosplenial cortex, parietal association cortex and posterior parietal cortex (either compared with their own baseline: Supplementary Table 8 or compared with the J60-treated shams: Supplementary Table 9). In addition, animals that received specific cholinergic MSDB stimulation consistently showed lower FC with prelimbic-infralimbic cortex than J60-treated sham rats (Supplementary Table 7).

With retrosplenial cortex as seed, there was also a significant change over time in FC in J60-treated shams (FDR-corrected p=0.0459) (Supplementary Table 8). However, the time effect on FC with retrosplenial cortex in MSDB-stimulated rats was much more pronounced (FDR-corrected p<0.0001 for both specifically and nonspecifically stimulated rats). FC with retrosplenial cortex was decreased compared with baseline in J60-treated shams only at 40-50 min p.i. (adj. p=0.0056), while FC was decreased in MSDB-stimulated rats during all post-J60 time intervals (adj. p<0.0001 for all time intervals for both specifically and nonspecifically stimulated rats). Timepoint-by-timepoint comparison showed that MSDB-stimulated rats always had a much lower FC with retrosplenial cortex after J60 injection than J60-treated sham rats (10-20, 20-30, 30-40 min p.i.: cholDREADD/J60 and DREADD/J60: adj. p<0.0001; 40-50 min p.i.: cholDREADD/J60: adj. p=0.0007, DREADD/J60: adj. p=0.0002; 50-60 min p.i.: cholDREADD/J60: adj. p=0.0008, DREADD/J60: adj. p<0.0001).

In agreement with the ROI-based FC analysis, animals receiving specific cholinergic MSDB stimulation showed a decreased FC after J60 injection with seeds from the hippocampus: the posterior hippocampus and dentate gyrus (compared with their own baseline: Supplementary Table 8 and compared with the J60-treated shams: Supplementary Table 9). These rats also showed consistently lower FC with anterior hippocampus than J60-treated sham rats (Supplementary Table 7). Also in agreement with the ROI-based FC analysis, animals receiving nonspecific neuronal MSDB stimulation had an increased FC with posterior hippocampus at 50-60 min p.i. compared with baseline (adj. p=0.0223).

Finally, this analysis revealed a significant effect of time on the FC with the nucleus accumbens in rats that received specific cholinergic MSDB stimulation (FDR-corrected p=0.0036), an effect that was absent in all other groups (FDR-corrected p>0.05) (Fig.5, Supplementary Table 8). More precisely, animals receiving specific cholinergic MSDB stimulation had a significantly higher FC with the nucleus accumbens during all post-J60 intervals (except the last) compared with baseline: 10-20 min p.i.: adj. p=0.0017, 20-30 min: adj. p=0.0015, 30-40 min: adj. p=0.0057, 40-50 min: adj. p=0.0009, and 50-60 min p.i.: adj. p=0.1710.

**Figure 5.**
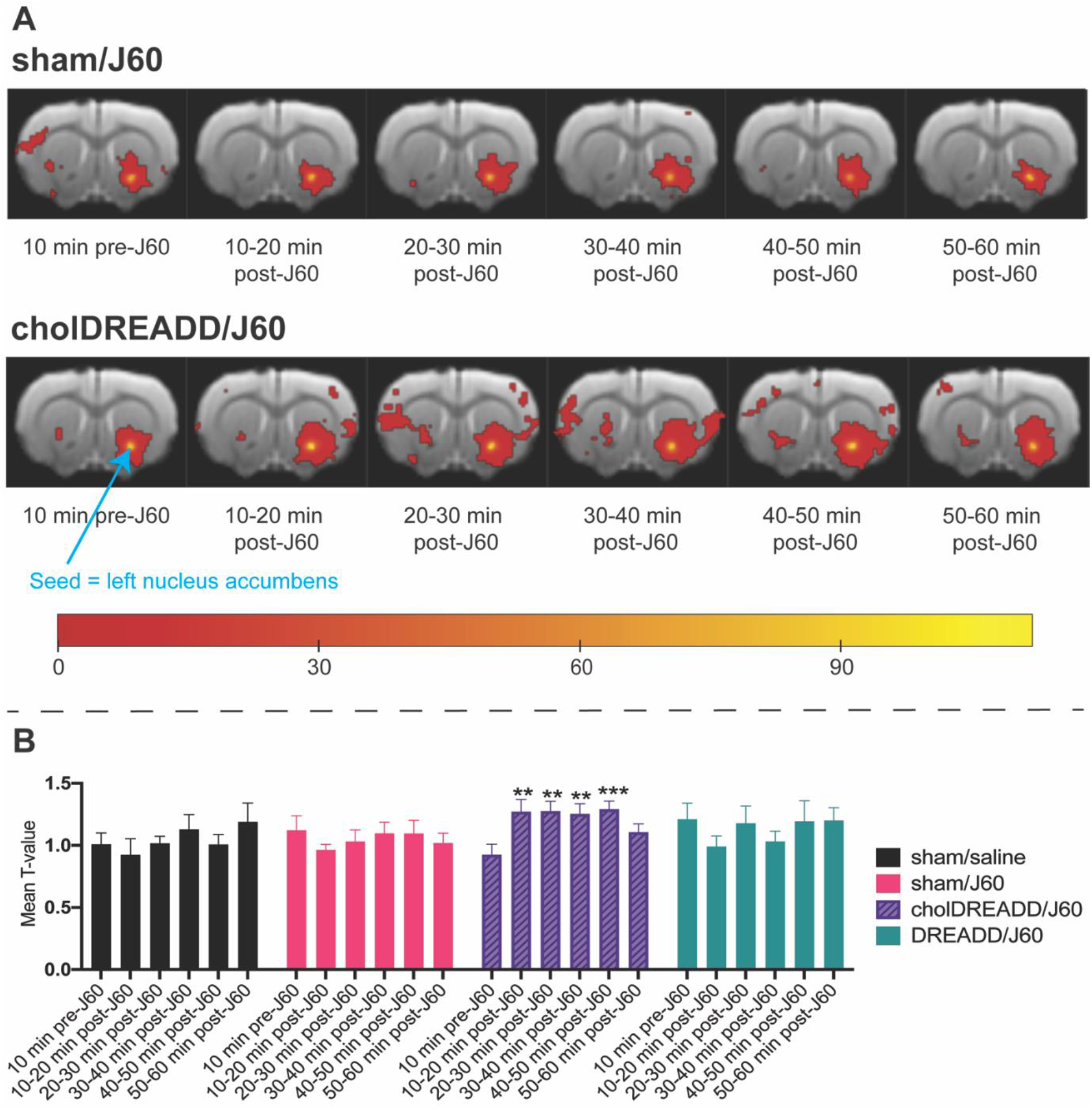
Seed-based functional connectivity (FC) analysis of the pharmacological fMRI data with the DREADD agonist J60 (or saline) as acute pharmacological challenge and with the left nucleus accumbens as seed. A. Group-averaged statistical seed-based FC maps with left nucleus accumbens as seed, indicated by the light blue arrow (one-sample t-tests, uncorrected, p≤0.001, minimal cluster size = 10 voxels). While FC with the left nucleus accumbens remains quite stable in J60-treated sham rats (sham/J60) over time, specific cholinergic medial septum-diagonal band of Broca (MSDB) stimulation (cholDREADD/J60) resulted in increased FC between the left nucleus accumbens and other parts of the salience-like network: the contralateral nucleus accumbens, ipsi- and contralateral ventral caudate putamen and insular cortex. The colour scale indicates T-values. B. Mean T-value within the summed mask based on the group-averaged statistical seed-based FC maps with left nucleus accumbens as seed (summed across all groups and all time intervals). Specific cholinergic MSDB stimulation (cholDREADD/J60, purple) resulted in increased FC with the left nucleus accumbens between 10 and 50 min post-J60-injection versus baseline. Sham/saline: n = 12, sham/J60: n = 14, cholDREADD/J60: n = 27, DREADD/J60: n = 14. **p≤0.01, ***p≤0.001. Only the significant changes within a group versus pre-J60 baseline are indicated on this graph.

#### The combined effect of radial arm maze memory training and chronic food restriction with or without chronic MSDB stimulation did not have an observable impact on resting-state networks

Resting-state FC in the five investigated network modules (anterior DMLN, posterior DMLN, hippocampus, LCN and SN) and resting-state FC with the 13 investigated seeds (OF, PrL-IL, Ant Cg, Post Cg, RS, PtA, PtP, Ant Hc, Post Hc, DG, Acb, CPu and Ins) did not significantly change over time, nor was there any difference between the different groups or a different effect of time in the different groups (FDR-corrected main time effects, main group effects and group*time interactions: p>0.05) (Fig.6, Supplementary Table 10). Hence, we can conclude that chronic MSDB stimulation (daily stimulation during memory training) had no observable effect on resting-state FC, since no difference was observed between MSDB-stimulated rats and sham rats during the post-acquisition training and post-reversal training resting-state fMRI scans. Since there were no FC differences between the pre-learning, post-acquisition and post-reversal resting-state fMRI scans, we can conclude that neither the combined effect of radial arm maze memory training, chronic food restriction and chronic MSDB stimulation (in MSDB-stimulated groups) nor the combined effect of radial arm maze memory training and chronic food restriction (in sham groups) had any observable effect on resting-state FC.

**Figure 6.**
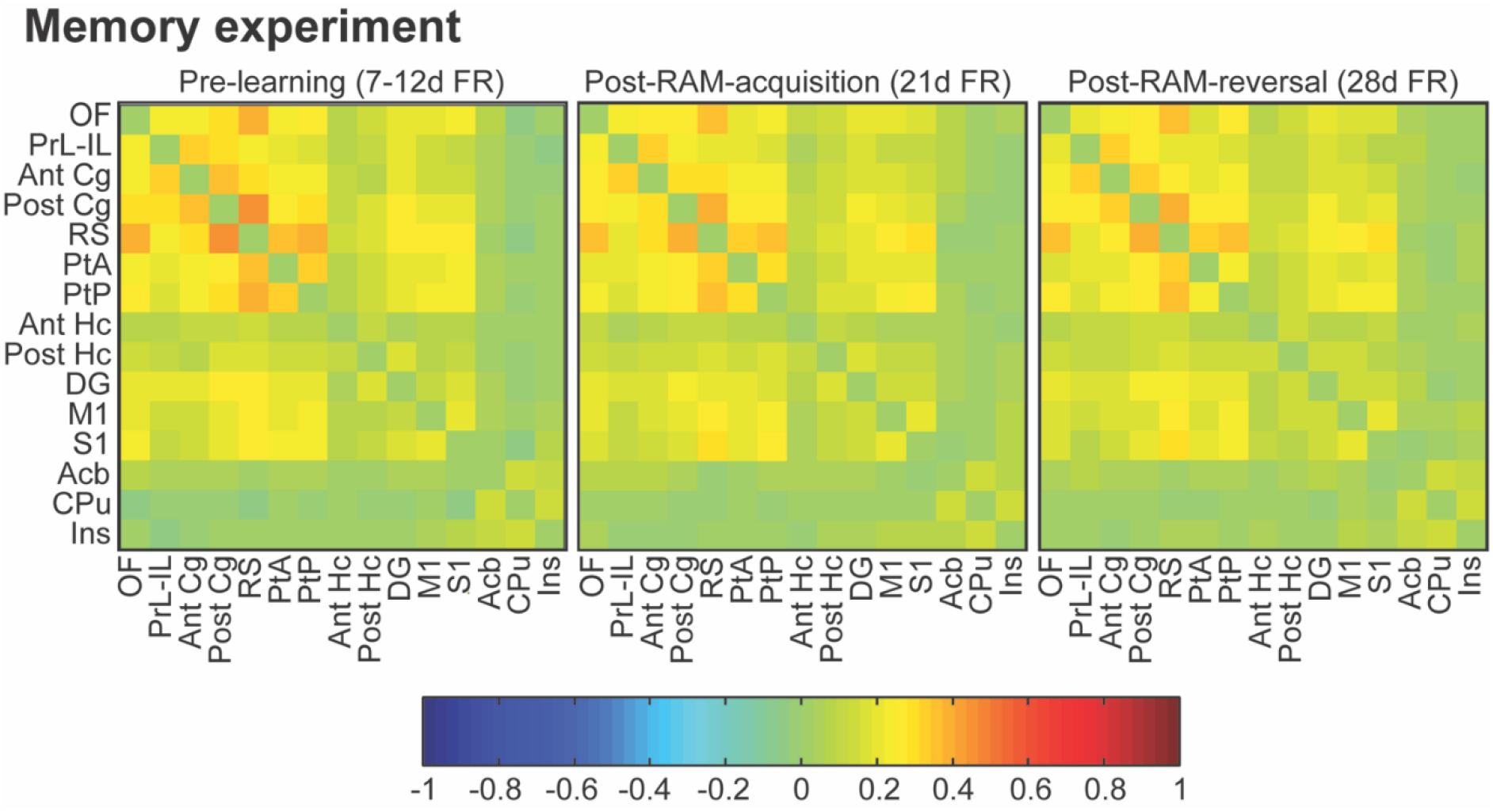
Region of interest (ROI)-based functional connectivity (FC) analysis of the resting-state fMRI data from the memory experiment. At the moment of the pre-learning scan, rats were food-restricted for at least 7 days and max. 12 days. During the post-radial arm maze (RAM)-acquisition training scan rats were food-restricted for 21 days and during the post-RAM-reversal training scan for 28 days. Group-averaged z-transformed FC matrices with the following ROIs from top to bottom (and from left to right) are presented: orbitofrontal cortex (OF), prelimbic and infralimbic cortices (PrL-IL), anterior cingulate cortex (Ant Cg), posterior cingulate cortex (Post Cg), retrosplenial cortex (RS), parietal association cortex (PtA), posterior parietal cortex (PtP), anterior hippocampus proper (Ant Hc), posterior hippocampus proper (Post Hc), dentate gyrus (DG), primary motor cortex (M1), primary somatosensory cortex (S1), nucleus accumbens (Acb), ventral caudate putamen (CPu) and insular cortex (Ins). Resting-state FC remained rather stable over time in the memory experiment, i.e. in rats that were both food-restricted for variable periods of time and during different stages of learning. The colour scale indicates the z-transformed correlation values. Z-transformed FC matrices are the average of all animals from all groups except the DREADD/J60/during group: sham/saline/during: n = 10, sham/J60/during: n = 9, cholDREADD/J60/during: n = 9, cholDREADD/J60/after: n = 10.

#### Chronic food restriction alone altered resting-state functional connectivity

Resting-state FC in the five investigated network modules (anterior DMLN, posterior DMLN, hippocampus, LCN and SN) did not significantly alter over time, though strong trends towards significance were observed for the anterior and posterior DMLN (FDR-corrected p=0.0515 for both) (Fig.7, Supplementary Table 11). Further investigation of these two network modules revealed a significant increase in FC at 28 days of food restriction compared with the pre-food restriction baseline (adj. p=0.0037 and adj. p=0.0031 for respectively anterior and posterior DMLN). However, there were no significant differences when comparing the FC with the 7-day food restriction timepoint (adj. p>0.05). We present the post-hoc analysis in two ways: i) comparing all timepoints with the pre-food restriction baseline to assess the effect of chronic food restriction on resting-state FC and ii) comparing all timepoints with the 7-day food restriction timepoint as this mimics most closely the baseline condition of the pre-learning fMRI scan. Hence, we can draw a better comparison with the resting-state fMRI data from the memory experiment.

**Figure 7.**
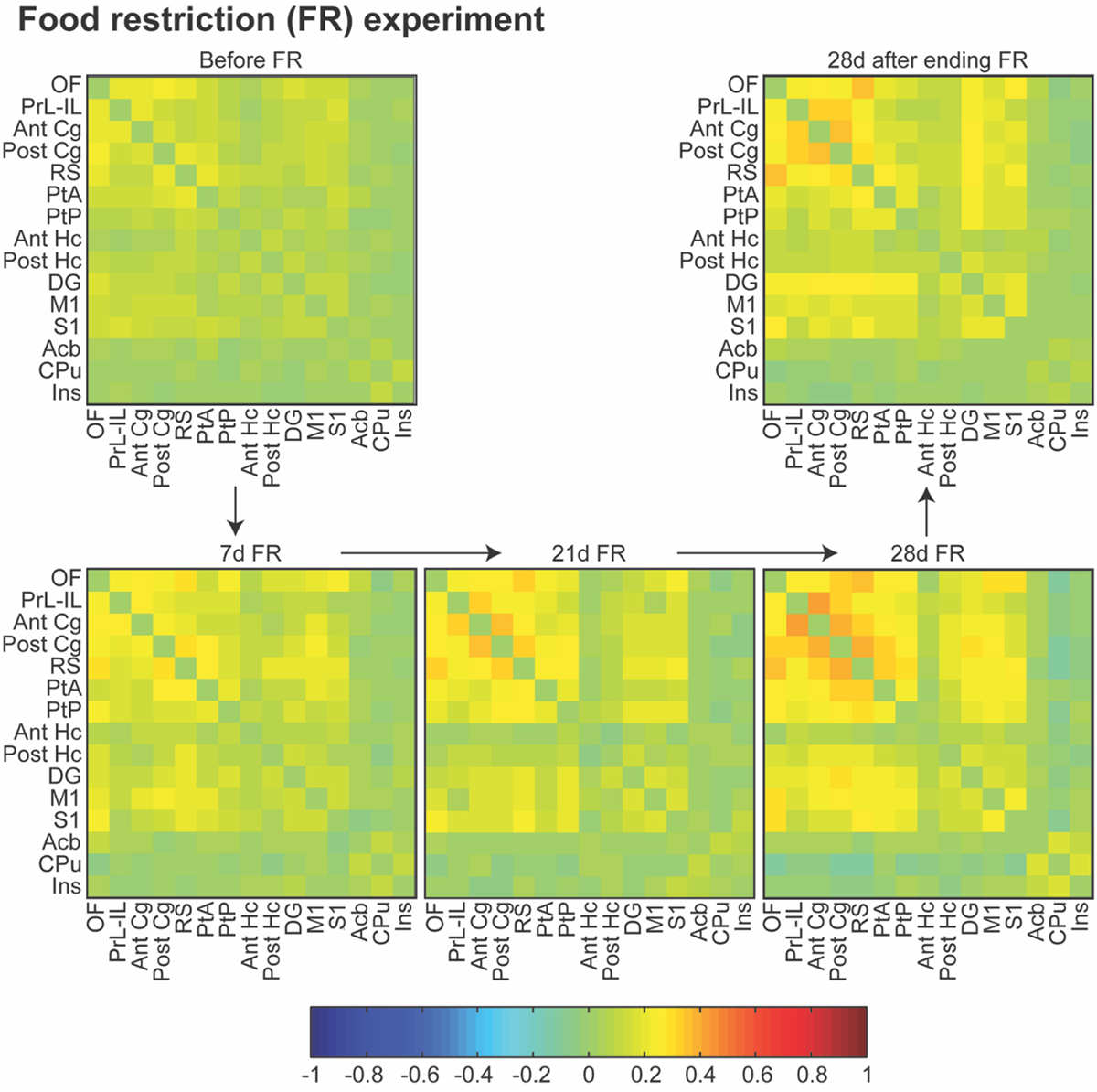
Region of interest (ROI)-based functional connectivity (FC) analysis of the resting-state fMRI data from the food restriction experiment. Rats were initially not food-restricted, then food-restricted for respectively 7 days, 21 days and 28 days and finally a scan was acquired when rats had not been food-restricted anymore for 28 days. Group-averaged z-transformed FC matrices with the following ROIs from top to bottom (and from left to right) are presented: orbitofrontal cortex (OF), prelimbic and infralimbic cortices (PrL-IL), anterior cingulate cortex (Ant Cg), posterior cingulate cortex (Post Cg), retrosplenial cortex (RS), parietal association cortex (PtA), posterior parietal cortex (PtP), anterior hippocampus proper (Ant Hc), posterior hippocampus proper (Post Hc), dentate gyrus (DG), primary motor cortex (M1), primary somatosensory cortex (S1), nucleus accumbens (Acb), ventral caudate putamen (CPu) and insular cortex (Ins). Food restriction in itself had a big effect on resting-state FC: FC increased with every additional week of food restriction and was still elevated four weeks after ending the four-week food restriction. The colour scale indicates the z-transformed correlation values. Z-transformed FC matrices are the average of twelve animals.

The seed-based FC analysis revealed a significant effect of time on FC with all investigated seeds except the anterior hippocampus (Supplementary Table 11). Overall, resting-state FC became higher the longer rats had been food-restricted, with the earliest significant changes versus the pre-food restriction baseline detectable at 21 days of food restriction (OF: adj. p=0.0405, Ant Cg: adj. p=0.0133, Post Cg: adj. p=0.0121, RS: adj. p=0.0064, and PtP: adj. p=0.0053). At 28 days of food restriction, there was increased FC compared with the pre-food restriction baseline for all seeds (OF, PrL-IL, Ant Cg, Post Cg, RS, PtA, PtP, DG and CPu: all adj. p<0.0001; Post Hc, Acb and Ins: all adj. p=0.0002). Surprisingly, 28 days after ending the food restriction, FC was still significantly increased (though less pronounced than at 28 days of food restriction) for OF (adj. p=0.0084), PrL-IL (adj. p=0.0126), Ant Cg (adj. p=0.0110), Post Cg (adj. p=0.0179), PtA (adj. p=0.0297), DG (adj. p=0.0047), CPu (adj. p=0.0083) and Ins (adj. p=0.0396) compared with the pre-food restriction baseline. Compared with the 7-day food restriction timepoint, significantly higher FC could only be observed at 28 days of food restriction, for all seeds except the orbitofrontal cortex (PrL-IL: adj. p=0.0049, Ant Cg: adj. p=0.0073, Post Cg: adj. p=0.0050, RS: adj. p=0.0237, PtA: adj. p=0.0007, PtP: adj. p=0.0006, Post Hc: adj. p=0.0160, DG: adj. p=0.0012, Acb: adj. p=0.0325, CPu: adj. p=0.0018, Ins: adj. p=0.0267).

Hence, we can conclude that chronic food restriction alone had a pronounced effect on resting-state FC. The effect became bigger the longer rats were food-restricted and had a long-lasting effect. Twenty-eight days after a 28-day period of food restriction, the FC was still increased compared with the pre-food restriction baseline. Even when comparing data with the 7-day food restriction timepoint, increased FC could be observed at 28 days of food restriction for almost all investigated seeds.

## Discussion

In summary, we observed that DREADD-based selective cholinergic MSDB stimulation initiated 15 min before radial arm maze training resulted in improved spatial learning/reference memory during the acquisition phase of the task but did not affect spatial working memory nor reversal learning. Selective cholinergic MSDB stimulation initiated shortly (on average (22 ± 1) min) after terminating radial arm maze training and approximately 15 min before the animals’ resting (light) phase started, had no effect on learning and memory. In addition, we observed that DREADD-based nonselective MSDB stimulation initiated 15 min before radial arm maze training resulted in aberrant behaviour in the majority of animals, most importantly characterised by a loss of appetite. J60-treated sham rats had a worse working memory than saline-treated sham rats during the reversal phase of the radial arm maze training, suggesting an adverse effect of 0.1 mg/kg J60 when used chronically. Both selective cholinergic and nonselective neuronal MSDB stimulation caused an acutely decreased functional connectivity in the default mode-like network (DMLN). Selective cholinergic MSDB stimulation resulted in increased functional connectivity with the nucleus accumbens and nonselective MSDB stimulation led to increased intrahippocampal functional connectivity. The combined effect of radial arm maze training and chronic food restriction with or without chronic MSDB stimulation caused no observable change in resting-state networks (RSNs). Chronic food restriction in itself had a marked effect on RSNs.

### Selective cholinergic neuronal MSDB stimulation during training, but not after training, improved spatial reference memory during acquisition in the radial arm maze

Selective cholinergic MSDB stimulation initiated 15 min before memory training started, improved acquisition of the spatial learning task. Most MSDB studies that showed improved cognition initiated MSDB stimulation immediately before the behavioural task started^10, 13, 14, 16, 17, 18, 19, 26, 27, 28^. Selective cholinergic MSDB stimulation initiated shortly (on average (22 ± 1) min) after the memory training ended, and approximately 15 min before the animals’ resting phase (light phase) started, did not improve spatial learning nor did it impair it. We hypothesised that cholinergic stimulation during slow-wave sleep would interfere with the memory consolidation during this sleep stage, which requires low hippocampal acetylcholine levels, and result in impaired declarative memory, as shown in human subjects^39^. However, cholinergic stimulation shortly after the memory training (during active wakefulness) and during REM sleep may have had beneficial effects, resulting eventually in a net effect of neither impaired nor improved spatial learning. Some studies that stimulated the MSDB shortly after the memory training have observed a beneficial effect on memory^12, 21, 23, 24, 25^. One of these studies observed that higher hippocampal cholinergic activity was correlated with lower consolidation capacities and suggested that the post-training electrical MSDB stimulation improved memory through non-cholinergic elements and that septohippocampal cholinergic activation is more likely to facilitate memory processes prior to consolidation^23^. In our study, a positive effect of post-training cholinergic MSDB stimulation on memory processes during active wakefulness and REM sleep, and a negative effect on the slow-wave sleep-related consolidation may have resulted in a net zero effect on memory. Differential effects of post-training MSDB stimulation between studies may be explained by a different timing of the MSDB stimulation during the day (light/dark phase) and thus different stages of the sleep-wake cycle that are affected by the MSDB stimulation.

Our study represents the first study where selective cholinergic MSDB stimulation resulted in improved learning and (reference) memory. Very recently, two other studies have investigated the effect of selective cholinergic MSDB stimulation on memory^46, 47^. Both of these studies used optogenetics to stimulate cholinergic MSDB neurons at a very specific time during the behavioural task.

Jarzebowski et al. stimulated septal cholinergic neurons at the goal zone in an appetitive Y-maze task, which slowed spatial learning. This coincided with a significant reduction in sharp wave ripple (SWR) incidence in hippocampal CA1 at the goal zone in rewarded trials, where SWRs most often occurred^46^. SWRs are an important oscillatory pattern in the hippocampus that are closely linked with (spatial) learning and memory. When a rodent follows a spatial trajectory, place cells are sequentially activated. During SWRs, this sequence of place cells is reactivated or replayed in a time-compressed manner in either a forward or backward direction. This event is thought to underlie memory consolidation. SWRs occur during slow-wave sleep, long periods of awake immobility and brief pauses in exploration. Hence, SWRs occur during stages of the sleep-wake cycle with low hippocampal acetylcholine levels (slow-wave sleep and quiet wakefulness) and are rare during stages with high hippocampal cholinergic tonus (active wakefulness and REM sleep)^48^. Interrupting SWRs has been shown to result in impaired spatial memory^49, 50, 51^. Selective optogenetic activation of septal cholinergic neurons has been shown to completely suppress SWRs in behaving and anaesthetised animals^52, 53^. Hence, it is unsurprising that optogenetic septal cholinergic stimulation at the goal zone reduced SWR incidence and impaired spatial memory. Cholinergic stimulation during navigation had no effect on spatial learning and animals stimulated throughout the maze (navigation + goal zones) had an intermediary learning curve^46^. It is surprising that we observed improved spatial learning in our rats with DREADD-mediated cholinergic MSDB stimulation throughout the maze, while Jarzebowski et al. observed a slight impairment of spatial learning with optogenetics-mediated cholinergic MSDB stimulation throughout the maze in their mice. Apart from different techniques and different species used in our studies, another difference exists, which could potentially explain the differential response. In our study, cholinergic MSDB neurons were activated for a longer time. They were already activated shortly before the start of the first behavioural trial, as well as between trials and for some time after the last behavioural trial. This is inherent to the technique that we used, since DREADDs lack the temporal precision of optogenetics and deep brain stimulation.

Zhang et al. selectively stimulated septal cholinergic neurons during the delay between choices in a spontaneous alternation task, when the hippocampal cholinergic signal was normally lowest and the SWR incidence highest. Optogenetic cholinergic activation caused an increase in hippocampal cholinergic signal, abolished SWRs, and impaired choice performance in this working memory task in mice^47^.

Though these studies show that one can interfere with memory by increasing acetylcholine signalling specifically at moments when the hippocampal acetylcholine is usually low and SWR incidence high, our study indicates that increasing cholinergic activity throughout the entire behavioural test, as well as shortly before the first trial, in between trials and after the last trial has a beneficial effect on spatial learning. In addition, we have also performed electrophysiological experiments and observed alterations in hippocampal oscillations after selective cholinergic neuronal MSDB stimulation via DREADDs (manuscript in preparation).

A few other studies have investigated the effects of selective activation or inhibition of septal cholinergic neurons on behaviour. hM3Dq DREADD-mediated activation of septal cholinergic neurons induced behaviours in mice associated with novelty or anxiety: freezing and thigmotaxis, and thus decreased exploration of the familiar environment^54^. hM4Di DREADD-mediated inhibition of septal cholinergic neurons had anxiolytic effects in mice and increased exploratory activities in an open field^55^. hM4Di DREADD-mediated inhibition of septal cholinergic neurons resulted in reduced prepulse inhibition, indicating the importance of septal cholinergic neurons in sensorimotor gating, the process of filtering out redundant or irrelevant stimuli^56^. In our study we did not observe freezing or decreased exploration in the radial arm maze after selective septal cholinergic activation.

### Selective cholinergic neuronal MSDB stimulation had no effect on spatial working memory or reversal learning in the radial arm maze

Selective cholinergic MSDB stimulation did not improve nor impair spatial working memory. Several MSDB stimulation studies, however, have observed improved spatial working memory^16, 17, 19, 28^. A possible explanation for the differential observation between these studies and our study is the difference in complexity of the working memory task. All of these studies made use of a T-maze rewarded alternation task, where there were only two arms to choose between. In our radial arm maze task, however, rats had to choose constantly between eight arms. Our task may simply have been too complicated to be able to detect an improvement in working memory.

Selective stimulation of cholinergic MSDB neurons did not improve or impair spatial reversal learning either. All our rats were capable of reversal learning as indicated by a significant effect of time during this learning phase. To our knowledge, no other studies have investigated the effect of MSDB stimulation on reversal learning. Flexible spatial learning has been shown to depend on the prefrontal cortex and hippocampus and their functional interactions^57^. It is possible that MSDB stimulation does not improve the interactions between hippocampus and prefrontal cortex, resulting thus in unaltered reversal learning.

### Nonselective neuronal MSDB stimulation during training resulted in aberrant behavior

Surprisingly, nonselective neuronal MSDB stimulation led to aberrant behaviour in most animals. The most astounding observation was a loss of appetite, despite the fact that animals were food-restricted. The animals neither ate the rewards in the maze nor all of the daily portion of regular food in their home cage following nonselective MSDB stimulation, leading them to be excluded from the study. We assume that the general loss of appetite underlay the loss of interest in the rewards and that the consequent loss of motivation to look for the rewards explains the significantly lower speed of the animals in the maze. During the habituation phase in the maze, these animals exhibited perfectly normal behaviour, including a normal interest in the rewards, and they always ate all of their daily allotted portion of food. Our study is the first study to use DREADDs for nonselective MSDB stimulation. Nonselective MSDB stimulation with optogenetics or deep brain stimulation has not been reported to result in such behaviour as we observed using DREADDs. This discrepancy may be explained by inherent differences between DREADDs on the one hand and optogenetics and deep brain stimulation on the other hand. Pulsed stimulation at a certain frequency, as is given with optogenetics and deep brain stimulation, may have a different effect than constant stimulation of MSDB neurons by binding of the DREADD agonist to the DREADD receptors. A study by Durkin in 1992 reported a similar observation upon intraseptal injection of the GABAA agonist muscimol or the GABAA antagonist bicuculline in mice during radial arm maze testing: subjects often stopped in front of the pellet tray for long periods without eating the reward and sometimes left the arm without having consumed the reward, a phenomenon they had never before observed in food-deprived mice during radial arm maze testing^58^. More recently, it has been shown that selective stimulation of glutamatergic neurons in the MSDB using DREADDs or optogenetics results in a loss of appetite in mice, probably by activating downstream neurons in the paraventricular hypothalamus^59^. Hence, we expect that stimulation of the glutamatergic MSDB neurons in our nonselective stimulation approach underlay the loss of appetite in our rats.

Some rats exhibited a pronounced increase in water consumption following nonselective MSDB stimulation. MSDB lesions in rats and sheep have been shown to cause a pronounced increase in water intake, indicating a role of the MSDB in drinking behaviour^60, 61^. We have no explanation for the fact that both MSDB stimulation in our study and MSDB lesions caused an increase in water intake. Liao and Yeh observed that only posterior lesions and not anterior lesions of the medial septum in rats caused an increase in water intake^60^. We aimed to transfect the middle of the medial septum with DREADDs in our study, but inter-subject variability can have caused some transfections to be more posterior, which could explain that only some of the animals had a marked increase in water intake.

### J60-treated sham rats had a worse spatial working memory during reversal learning in the radial arm maze than saline-treated sham rats

During reversal learning, in the third week of daily J60 injections, J60-treated sham rats had a worse working memory than saline-treated sham rats in the radial arm maze. This suggests that chronic use of 0.1 mg/kg J60 had an adverse effect. Our study is the first study to use the novel DREADD agonist J60 in a chronic set-up. Previously published studies that made use of J60 only administered a few doses^36, 37, 38^. Our study underlines the importance of including a DREADD-free control group that is treated with the DREADD agonist to account for adverse effects of the DREADD agonist itself. This is the control group that we used as a reference to compare all other groups with. However, one needs to keep in mind that the adverse effects of a DREADD agonist may only occur in DREADD-free animals. It is possible that when DREADDs are present, the DREADD ligand will only bind to the DREADDs for which it has the highest affinity and not to endogenous receptors for which it has a lower affinity. In DREADD-free animals the DREADD ligand will bind to endogenous receptors in the absence of another target and can thus cause adverse effects that may not occur when DREADDs are expressed in the animal.

### Pharmacological functional connectivity fMRI reveals a differential impact of selective cholinergic and nonselective neuronal MSDB stimulation on resting-state networks (RSNs)

Pharmacological functional connectivity fMRI with the DREADD agonist J60 or its vehicle saline revealed that acute administration of a single dose of J60 or a saline injection did not affect RSNs in DREADD-free animals.

In DREADD-expressing animals, both selective cholinergic and nonselective neuronal MSDB stimulation induced an acute decrease in functional connectivity within the default mode-like network (DMLN). It has been shown that the basal forebrain contributes to DMLN regulation^62^ and our research group has previously demonstrated that DREADD-mediated selective cholinergic stimulation of the right basal forebrain (horizontal limb of the diagonal band of Broca, substantia innominata, and nucleus basalis of Meynert) decreased functional connectivity in the DMLN^63^. The DMLN is suppressed during cognitive tasks to quieten internal activity and optimise externally directed cognitive function^64^. MSDB stimulation during our radial arm maze task may have enhanced DMLN suppression, thus improving the externally directed cognitive function, which may have underlain the enhanced performance in the radial arm maze observed in cholinergically MSDB stimulated rats.

Nonselective neuronal MSBD stimulation caused increased intrahippocampal functional connectivity (FC), while selective cholinergic MSDB stimulation did not. A relationship has been shown between BOLD fMRI signal and local field potentials in the theta band in the human hippocampal area^65^. Theta oscillations are closely linked to memory and navigation and the MSDB seems to act as the pacemaker of this hippocampal theta rhythm. Strong evidence indicates that GABAergic and glutamatergic MSDB neurons play an important role in the generation of theta rhythm, while cholinergic MSDB neurons play a less important role^66^. It is possible that the increased intrahippocampal FC observed following nonselective MSDB stimulation but not after selective cholinergic MSDB stimulation reflects a difference in the capacity of the two stimulation protocols to alter hippocampal theta rhythm^67^. Unfortunately, the memory of nonspecifically stimulated rats could not be investigated due to the above-mentioned loss of appetite.

Finally, selective cholinergic but not nonselective MSDB stimulation caused increased FC with the nucleus accumbens. The MSDB projects to the ventral subiculum (part of the hippocampal formation), which in its turn projects to the nucleus accumbens. From there it projects via the rostral ventral pallidum to the ventral tegmental area. MSDB activation has been shown to increase dopaminergic neuron population activity in the ventral tegmental area via this pathway, which depended selectively on cholinergic mechanisms in the ventral subiculum (most likely the cholinergic septohippocampal neurons)^68^. These dopaminergic neurons in the ventral tegmental area project to the nucleus accumbens, constituting the mesolimbic pathway or reward pathway. Release of dopamine into the nucleus accumbens regulates incentive salience (“wanting” a reward) and facilitates reinforcement and reward-related learning of associations between sensory cues and profitable motor responses^69, 70^. “Wanting” a reward can be dramatically intensified in reencounter if dopamine levels in the nucleus accumbens are elevated above normal by a drug like amphetamine for instance^69^. In our study, selective stimulation of cholinergic MSDB neurons may have increased dopaminergic neuron population activity in the ventral tegmental area and hence increased dopamine levels in the nucleus accumbens during the radial arm maze task, resulting in increased incentive salience (“wanting”), enhanced motivation and improved learning of the reward-based task.

### The combined effect of radial arm maze memory training and chronic food restriction with or without chronic MSDB stimulation did not have an observable impact on RSNs, but chronic food restriction in itself did have a marked effect on RSNs

The combination of radial arm maze learning and chronic food restriction with and without chronic MSDB stimulation did not have an observable effect on RSNs. However, chronic food restriction alone did have a significant effect on RSNs. FC increased over time with prolonged food restriction and remained elevated for a long time after ending the food restriction. This effect of chronic food restriction on FC may have masked the effects of radial arm maze training and chronic MSDB stimulation on RSNs. Acute food restriction (twelve hours of fasting) has also been shown to cause globally increased FC in mice^71^. A study in which rats were food-restricted for one week observed no significant effect on FC, though a trend towards lower FC in the food-restricted rats was reported^72^. We also did not observe a significant effect on FC after only one week of food restriction.

Since chronic food restriction caused the rats to be more agitated than we are used to, we gave rats a bit more isoflurane than usual during the positioning of the rat in the scanner bed.

This was kept constant for all rats during all scanning sessions at all timepoints, including the scanning session before food restriction in cohort 2. This could explain why we observed such low FC in the pre-food restriction scanning session. Normally, we scan rats without any food restriction with a lower dose of isoflurane during the set-up. Otherwise, the anaesthesia protocol was exactly the same as published before^45^.

### Conclusion and future perspectives

In conclusion, we compared the effects of a DREADD-mediated selective cholinergic and nonselective neuronal MSDB stimulation strategy on spatial learning and memory and RSNs in the brain. Selective cholinergic MSDB stimulation during, but not after, radial arm maze training improved spatial (acquisition) learning, possibly by enhanced DMLN suppression and increased functioning of the nucleus accumbens during task performance. Nonselective neuronal MSDB stimulation increased intrahippocampal FC, but also caused a loss of appetite. Because of this, learning and memory could not properly be investigated using the appetitive radial arm maze task.

Next, the selective cholinergic MSDB stimulation strategy should be tested in animal models with cholinergic and cognitive deficits to evaluate if the cognitive deficits can be rescued by stimulating the surviving cholinergic basal forebrain neurons. If successful, this could then be used as an alternative therapy to the acetylcholinesterase inhibitors, which are used in patients with Alzheimer’s disease and dementia with Lewy bodies. It would provide these patients with an alternative therapy without adverse effects. For patients with Alzheimer’s disease, the focus should be shifted to stimulation of the nucleus basalis of Meynert, since cholinergic neuronal loss is observed in this basal forebrain region, which is the source of cortical cholinergic innervation^73^. However, patients suffering from dementia with Lewy bodies could benefit from stimulation of both the nucleus basalis of Meynert and the MSDB, since cholinergic neuronal loss has been observed in both these regions in these patients^74^. Finally, it should be investigated whether chronic use of a lower dose of J60 than the one used here (0.1 mg/kg) does not lead to adverse effects or if another novel DREADD agonist, e.g. JHU37152 (“J52”) could be used instead without any adverse effects in a chronic setting^36^.

## Acknowledgments

Stephan Missault was a postdoctoral fellow of the Research Foundation Flanders (FWO) (12W1619N). This research was funded by the Research Foundation Flanders (FWO) (G048917N) and the University Research Fund of the University of Antwerp (Small Research Grant BOF FFB200054). The 7T PharmaScan MR system was purchased through Hercules foundation funding (Belgium) under the promotership of Prof. Annemie Van der Linden. The computational resources and services used in this work for making the EPI template and realigning the pharmacological functional connectivity fMRI data were provided by the HPC core facility CalcUA of the Universiteit Antwerpen, the VSC (Flemish Supercomputer Center), funded by the Hercules Foundation and the Flemish Government – department EWI.

## References

1. Khakpai F, Nasehi M, Haeri-Rohani A, Eidi A, Zarrindast MR. Septo-hippocampo-septal loop and memory formation. Basic Clin Neurosci 4, 5–23 (2013).

2. Gold PE. Acetylcholine modulation of neural systems involved in learning and memory. Neurobiol Learn Mem 80, 194–210 (2003).

3. Woolf NJ, Eckenstein F, Butcher LL. Cholinergic systems in the rat brain: I. projections to the limbic telencephalon. Brain Res Bull 13, 751–784 (1984).

4. Paul S, Jeon WK, Bizon JL, Han JS. Interaction of basal forebrain cholinergic neurons with the glucocorticoid system in stress regulation and cognitive impairment. Front Aging Neurosci 7, 43 (2015).

5. Davies P, Maloney AJ. Selective loss of central cholinergic neurons in Alzheimer’s disease. Lancet 2, 1403 (1976).

6. Whitehouse PJ, Price DL, Clark AW, Coyle JT, DeLong MR. Alzheimer disease: evidence for selective loss of cholinergic neurons in the nucleus basalis. Ann Neurol 10, 122–126 (1981).

7. Xu H, Garcia-Ptacek S, Jonsson L, Wimo A, Nordstrom P, Eriksdotter M. Long-term Effects of Cholinesterase Inhibitors on Cognitive Decline and Mortality. Neurology 96, e2220–e2230 (2021).

8. Ali TB, Schleret TR, Reilly BM, Chen WY, Abagyan R. Adverse Effects of Cholinesterase Inhibitors in Dementia, According to the Pharmacovigilance Databases of the United-States and Canada. PLoS One 10, e0144337 (2015).

9. Lee DJ, et al. Medial septal nucleus theta frequency deep brain stimulation improves spatial working memory after traumatic brain injury. J Neurotrauma 30, 131–139 (2013).

10. Lee DJ, et al. Septohippocampal Neuromodulation Improves Cognition after Traumatic Brain Injury. J Neurotrauma 32, 1822–1832 (2015).

11. Jeong DU, Lee JE, Lee SE, Chang WS, Kim SJ, Chang JW. Improvements in memory after medial septum stimulation are associated with changes in hippocampal cholinergic activity and neurogenesis. Biomed Res Int 2014, 568587 (2014).

12. Jeong DU, Lee J, Chang WS, Chang JW. Identifying the appropriate time for deep brain stimulation to achieve spatial memory improvement on the Morris water maze. BMC Neurosci 18, 29 (2017).

13. Izadi A, Pevzner A, Lee DJ, Ekstrom AD, Shahlaie K, Gurkoff GG. Medial septal stimulation increases seizure threshold and improves cognition in epileptic rats. Brain Stimul 12, 735–742 (2019).

14. Lee DJ, et al. Stimulation of the medial septum improves performance in spatial learning following pilocarpine-induced status epilepticus. Epilepsy Res 130, 53–63 (2017).

15. Wang Y, et al. Deep brain stimulation in the medial septum attenuates temporal lobe epilepsy via entrainment of hippocampal theta rhythm. CNS Neurosci Ther 27, 577–586 (2021).

16. Li L, et al. Activation and blockade of 5-HT6 receptor in the medial septum-diagonal band recover working memory in the hemiparkinsonian rats. Brain Res 1748, 147072 (2020).

17. Li LB, et al. Activation of serotonin2A receptors in the medial septum-diagonal band of Broca complex enhanced working memory in the hemiparkinsonian rats. Neuropharmacology 91, 23–33 (2015).

18. Sava S, Markus EJ. Activation of the medial septum reverses age-related hippocampal encoding deficits: a place field analysis. J Neurosci 28, 1841–1853 (2008).

19. Markowska AL, Olton DS, Givens B. Cholinergic manipulations in the medial septal area: age-related effects on working memory and hippocampal electrophysiology. J Neurosci 15, 2063–2073 (1995).

20. Etter G, van der Veldt S, Manseau F, Zarrinkoub I, Trillaud-Doppia E, Williams S. Optogenetic gamma stimulation rescues memory impairments in an Alzheimer’s disease mouse model. Nature Communications 10, (2019).

21. Wetzel W, Ott T, H AM. Post-training hippocampal rhythmic slow activity (“theta”) elicited by septal stimulation improves memory consolidation in rats. Behav Biol 21, 32–40 (1977).

22. Deupree D, Coppock W, Willer H. Pretraining septal driving of hippocampal rhythmic slow activity facilitates acquisition of visual discrimination. J Comp Physiol Psychol 96, 557–562 (1982).

23. Galey D, Destrade C, Jaffard R. Relationships between septo-hippocampal cholinergic activation and the improvement of long-term retention produced by medial septal electrical stimulation in two inbred strains of mice. Behav Brain Res 60, 183–189 (1994).

24. 24. Izquierdo I, da Cunha C, Rosat R, Jerusalinsky D, Ferreira MB, Medina JH. Neurotransmitter receptors involved in post-training memory processing by the amygdala, medial septum, and hippocampus of the rat. Behav Neural Biol 58, 16–26 (1992).

25. Izquierdo I, Medina JH, Jeriisalinsky D, Da Cunha C. Post-Training Memory Processing in Amygdala, Septum and Hippocampus: Role of Benzodiazepine/GABAA Receptors, and their Interaction with other Neurotransmitter Systems. Rev Neurosci 3, 11–24 (1992).

26. Yousefi B, Nasehi M, Khakpai F, Zarrindast MR. Possible interaction of cholinergic and GABAergic systems between MS and CA1 upon memory acquisition in rats. Behav Brain Res 235, 231–243 (2012).

27. Micheau J, Van Marrewijk B. Stimulation of 5-HT1A receptors by systemic or medial septum injection induces anxiogenic-like effects and facilitates acquisition of a spatial discrimination task in mice. Prog Neuropsychopharmacol Biol Psychiatry 23, 1113–1133 (1999).

28. Blumberg BJ, et al. Efficacy of nonselective optogenetic control of the medial septum over hippocampal oscillations: the influence of speed and implications for cognitive enhancement. Physiol Rep 4, (2016).

29. Sans-Dublanc A, Razzauti A, Desikan S, Pascual M, Monyer H, Sindreu C. Septal GABAergic inputs to CA1 govern contextual memory retrieval. Sci Adv 6, (2020).

30. Luttgen M, Ogren SO, Meister B. 5-HT1A receptor mRNA and immunoreactivity in the rat medial septum/diagonal band of Broca-relationships to GABAergic and cholinergic neurons. J Chem Neuroanat 29, 93–111 (2005).

31. Luttgen M, Ove Ogren S, Meister B. Chemical identity of 5-HT2A receptor immunoreactive neurons of the rat septal complex and dorsal hippocampus. Brain Res 1010, 156–165 (2004).

32. Abode-Iyamah KO, et al. Deep brain stimulation hardware-related infections: 10-year experience at a single institution. J Neurosurg, 1–10 (2018).

33. Gomez JL, et al. Chemogenetics revealed: DREADD occupancy and activation via converted clozapine. Science 357, 503–507 (2017).

34. MacLaren DA, et al. Clozapine N-Oxide Administration Produces Behavioral Effects in Long-Evans Rats: Implications for Designing DREADD Experiments. eNeuro 3, (2016).

35. Manvich DF, et al. The DREADD agonist clozapine N-oxide (CNO) is reverse-metabolized to clozapine and produces clozapine-like interoceptive stimulus effects in rats and mice. Sci Rep 8, 3840 (2018).

36. Bonaventura J, et al. High-potency ligands for DREADD imaging and activation in rodents and monkeys. Nat Commun 10, 4627 (2019).

37. Amorim MR, et al. The Effect of DREADD Activation of Leptin Receptor Positive Neurons in the Nucleus of the Solitary Tract on Sleep Disordered Breathing. Int J Mol Sci 22, (2021).

38. Fleury Curado T, et al. Designer Receptors Exclusively Activated by Designer Drugs Approach to Treatment of Sleep-disordered Breathing. Am J Respir Crit Care Med 203, 102–110 (2021).

39. Gais S, Born J. Low acetylcholine during slow-wave sleep is critical for declarative memory consolidation. Proc Natl Acad Sci U S A 101, 2140–2144 (2004).

40. Hasselmo ME. The role of acetylcholine in learning and memory. Curr Opin Neurobiol 16, 710–715 (2006).

41. Power AE. Slow-wave sleep, acetylcholine, and memory consolidation. Proc Natl Acad Sci U S A 101, 1795–1796 (2004).

42. Albert NB, Robertson EM, Mehta P, Miall RC. Resting state networks and memory consolidation. Commun Integr Biol 2, 530–532 (2009).

43. Shah D, Verhoye M, Van der Linden A, D’Hooge R. Acquisition of Spatial Search Strategies and Reversal Learning in the Morris Water Maze Depend on Disparate Brain Functional Connectivity in Mice. Cereb Cortex 29, 4519–4529 (2019).

44. Witten IB, et al. Recombinase-driver rat lines: tools, techniques, and optogenetic application to dopamine-mediated reinforcement. Neuron 72, 721–733 (2011).

45. Missault S, et al. Hypersynchronicity in the default mode-like network in a neurodevelopmental animal model with relevance for schizophrenia. Behav Brain Res 364, 303–316 (2019).

46. Jarzebowski P, Tang CS, Paulsen O, Hay YA. Impaired spatial learning and suppression of sharp wave ripples by cholinergic activation at the goal location. Elife 10, (2021).

47. Zhang Y, et al. Cholinergic suppression of hippocampal sharp-wave ripples impairs working memory. Proc Natl Acad Sci U S A 118, (2021).

48. Atherton LA, Dupret D, Mellor JR. Memory trace replay: the shaping of memory consolidation by neuromodulation. Trends Neurosci 38, 560–570 (2015).

49. Ego-Stengel V, Wilson MA. Disruption of ripple-associated hippocampal activity during rest impairs spatial learning in the rat. Hippocampus 20, 1–10 (2010).

50. Girardeau G, Benchenane K, Wiener SI, Buzsaki G, Zugaro MB. Selective suppression of hippocampal ripples impairs spatial memory. Nat Neurosci 12, 1222–1223 (2009).

51. Jadhav SP, Kemere C, German PW, Frank LM. Awake hippocampal sharp-wave ripples support spatial memory. Science 336, 1454–1458 (2012).

52. Vandecasteele M, et al. Optogenetic activation of septal cholinergic neurons suppresses sharp wave ripples and enhances theta oscillations in the hippocampus. Proc Natl Acad Sci U S A 111, 13535–13540 (2014).

53. Ma X, et al. The Firing of Theta State-Related Septal Cholinergic Neurons Disrupt Hippocampal Ripple Oscillations via Muscarinic Receptors. J Neurosci 40, 3591–3603 (2020).

54. Carpenter F, Burgess N, Barry C. Modulating medial septal cholinergic activity reduces medial entorhinal theta frequency without affecting speed or grid coding. Sci Rep 7, 14573 (2017).

55. Zhang Y, et al. Inhibiting medial septal cholinergic neurons with DREADD alleviated anxiety-like behaviors in mice. Neurosci Lett 638, 139–144 (2017).

56. Jin J, et al. Cholinergic Neurons of the Medial Septum Are Crucial for Sensorimotor Gating. J Neurosci 39, 5234–5242 (2019).

57. Avigan PD, Cammack K, Shapiro ML. Flexible spatial learning requires both the dorsal and ventral hippocampus and their functional interactions with the prefrontal cortex. Hippocampus 30, 733–744 (2020).

58. Durkin TP. GABAergic mediation of indirect transsynaptic control over basal and spatial memory testing-induced activation of septo-hippocampal cholinergic activity in mice. Behav Brain Res 50, 155–165 (1992).

59. Sweeney P, Li C, Yang Y. Appetite suppressive role of medial septal glutamatergic neurons. Proc Natl Acad Sci U S A 114, 13816–13821 (2017).

60. Liao RM, Yeh CC. Influences on water intake in the rat after lesions of the septal subareas. Proc Natl Sci Counc Repub China B 24, 26–32 (2000).

61. Smardencas A, Denton DA, McKinley MJ. Hyperdipsia in sheep bearing lesions in the medial septal nucleus. Brain Res 1752, 147223 (2021).

62. Nair J, Klaassen AL, Arato J, Vyssotski AL, Harvey M, Rainer G. Basal forebrain contributes to default mode network regulation. Proc Natl Acad Sci U S A 115, 1352–1357 (2018).

63. Peeters LM, van den Berg M, Hinz R, Majumdar G, Pintelon I, Keliris GA. Cholinergic Modulation of the Default Mode Like Network in Rats. iScience 23, 101455 (2020).

64. Anticevic A, Cole MW, Murray JD, Corlett PR, Wang XJ, Krystal JH. The role of default network deactivation in cognition and disease. Trends Cogn Sci 16, 584–592 (2012).

65. Ekstrom A, Suthana N, Millett D, Fried I, Bookheimer S. Correlation between BOLD fMRI and theta-band local field potentials in the human hippocampal area. J Neurophysiol 101, 2668–2678 (2009).

66. Nunez A, Buno W. The Theta Rhythm of the Hippocampus: From Neuronal and Circuit Mechanisms to Behavior. Front Cell Neurosci 15, 649262 (2021).

67. Mamad O, McNamara HM, Reilly RB, Tsanov M. Medial septum regulates the hippocampal spatial representation. Front Behav Neurosci 9, 166 (2015).

68. Bortz DM, Grace AA. Medial septum differentially regulates dopamine neuron activity in the rat ventral tegmental area and substantia nigra via distinct pathways. Neuropsychopharmacology 43, 2093–2100 (2018).

69. Berridge KC. From prediction error to incentive salience: mesolimbic computation of reward motivation. Eur J Neurosci 35, 1124–1143 (2012).

70. Gale JT, Shields DC, Ishizawa Y, Eskandar EN. Reward and reinforcement activity in the nucleus accumbens during learning. Front Behav Neurosci 8, 114 (2014).

71. Tsurugizawa T, Djemai B, Zalesky A. The impact of fasting on resting state brain networks in mice. Sci Rep 9, 2976 (2019).

72. Roelofs TJM, Straathof M, van der Toorn A, Otte WM, Adan RAH, Dijkhuizen RM. Diet as connecting factor: Functional brain connectivity in relation to food intake and sucrose tasting, assessed with resting-state functional MRI in rats. J Neurosci Res, (2019).

73. Liu AK, Chang RC, Pearce RK, Gentleman SM. Nucleus basalis of Meynert revisited: anatomy, history and differential involvement in Alzheimer’s and Parkinson’s disease. Acta Neuropathol 129, 527–540 (2015).

74. Fujishiro H, Umegaki H, Isojima D, Akatsu H, Iguchi A, Kosaka K. Depletion of cholinergic neurons in the nucleus of the medial septum and the vertical limb of the diagonal band in dementia with Lewy bodies. Acta Neuropathol 111, 109–114 (2006).

