## Supplementary data for "Selective cholinergic stimulation of the medial septum-diagonal band of Broca via DREADDs improves spatial learning in healthy rats"

*Supplementary Table 1. Statistical analysis of the radial arm maze memory training: main effects. *p≤0.05, **p≤0.01, ***p≤0.001*

| Training phase | Parameter | P-value group effect | P-value time effect | P-value group*time interaction |
|---|---|---|---|---|
| Acquisition training | Total duration | **<0.0001*** ** | **<0.0001*** ** | 0.9263 |
| | Total distance | 0.2744 | **<0.0001*** ** | 0.2847 |
| | Mean speed | **0.0024** ** | **0.0051** ** | 0.7216 |
| | % reference memory errors | **0.0068** ** | **<0.0001*** ** | 0.3218 |
| | % time spent in non-baited arms | 0.0798 | **<0.0001*** ** | 0.3024 |
| | % distance travelled in non-baited arms | **0.0103* ** | **<0.0001*** ** | 0.1709 |
| | Latency to entry in first non-baited arm | **0.0346* ** | **<0.0001*** ** | 0.1030 |
| | Latency to entry in first baited arm | **0.0074** ** | **<0.0001*** ** | 0.9157 |
| | % working memory errors | **0.0247* ** | **<0.0001*** ** | 0.8571 |
| Reversal training | Total duration | 0.3717 | **<0.0001*** ** | 0.1392 |
| | Total distance | 0.0190* | <0.0001*** | **0.0216* ** |
| | Mean speed | 0.1855 | **<0.0001*** ** | 0.1246 |
| | % reference memory errors | 0.1256 | **<0.0001*** ** | 0.3215 |
| | % time spent in non-baited arms | **0.0148* ** | **<0.0001*** ** | 0.5239 |
| | % distance travelled in non-baited arms | 0.5386 | **<0.0001*** ** | 0.3515 |
| | Latency to entry in first non-baited arm | 0.4857 | **<0.0001*** ** | 0.7791 |
| | Latency to entry in first baited arm | 0.3514 | **0.0142* ** | 0.5952 |
| | % working memory errors | **0.0229* ** | **<0.0001*** ** | 0.1345 |

*Supplementary Table 2. Statistical analysis of the radial arm maze acquisition training: post-hoc analysis of the significant group effects from Supplementary Table 1. *p≤0.05, **p≤0.01, ***p≤0.001*

| **Training phase** | **Parameter** | **Post-hoc analysis of group effect: Dunnett's multiple comparisons with control test with sham/J60/during rats as control group: adjusted p-value + direction of effect vs. sham/J60/during** | |
|---|---|---|---|
| Acquisition training | Total duration | sham/saline/during | 0.9713 |
| | | cholDREADD/J60/during | 0.9760 |
| | | cholDREADD/J60/after | 0.9129 |
| | | **DREADD/J60/during** | **0.0003*** (>)** |
| | Mean speed | sham/saline/during | 0.8671 |
| | | cholDREADD/J60/during | 0.5804 |
| | | cholDREADD/J60/after | 0.9999 |
| | | **DREADD/J60/during** | **0.0011** (<)** |
| | % reference memory errors | sham/saline/during | 0.4950 |
| | | **cholDREADD/J60/during** | **0.0150* (<)** |
| | | cholDREADD/J60/after | 0.5020 |
| | | DREADD/J60/during | 0.5509 |
| | % distance travelled in non-baited arms | sham/saline/during | 0.4927 |
| | | **cholDREADD/J60/during** | **0.0129* (<)** |
| | | cholDREADD/J60/after | 0.1202 |
| | | DREADD/J60/during | 0.9336 |
| | Latency to entry in first non-baited arm | sham/saline/during | 0.7955 |
| | | **cholDREADD/J60/during** | **0.0199* (>)** |
| | | cholDREADD/J60/after | 0.9995 |
| | | DREADD/J60/during | 0.9997 |
| | Latency to entry in first baited arm | sham/saline/during | 1.0000 |
| | | cholDREADD/J60/during | 0.9955 |
| | | cholDREADD/J60/after | 0.9998 |
| | | **DREADD/J60/during** | **0.0106* (>)** |
| | % working memory errors | sham/saline/during | 0.4323 |
| | | cholDREADD/J60/during | 0.5307 |
| | | cholDREADD/J60/after | 0.7166 |
| | | DREADD/J60/during | 0.2142 |

*Supplementary Table 3. Statistical analysis of the radial arm maze reversal training: post-hoc analysis of the significant group*time interaction and group effects from Supplementary Table 1. *p≤0.05, **p≤0.01, ***p≤0.001*

| **Training phase** | **Parameter** | **Post-hoc analysis of group*time interaction: analysis per group**<br>**Time effects per group: FDR-corrected p-values** | |
|---|---|---|---|
| Reversal training | Total distance | **sham/saline/during** | **0.0003*** ** |
| | | **sham/J60/during** | **0.0001** ** |
| | | **cholDREADD/J60/during** | **0.0111*** |
| | | **cholDREADD/J60/after** | **0.0013** ** |
| | | DREADD/J60/during | 0.0804 |
| | | **Post-hoc analysis of group effect: Dunnett's multiple comparisons with control test with sham/J60/during rats as control group: adjusted p-value + direction of effect vs. sham/J60/during** | |
| | % time spent in non-baited arms | sham/saline/during | 0.9614 |
| | | cholDREADD/J60/during | 0.8848 |
| | | cholDREADD/J60/after | 1.0000 |
| | | **DREADD/J60/during** | **0.0142* (>)** |
| | % working memory errors | **sham/saline/during** | **0.0129* (<)** |
| | | cholDREADD/J60/during | 0.0547 |
| | | cholDREADD/J60/after | 0.1580 |
| | | DREADD/J60/during | 1.0000 |

*Supplementary Table 4. Statistical analysis of the ROI-based functional connectivity analysis of the pharmacological functional connectivity fMRI data with J60 or saline: main effects and post-hoc analysis of the significant group effect. DMLN = default mode-like network, LCN = lateral cortical network, SN = salience-like network. *p≤0.05, **p≤0.01, ***p≤0.001*

| Network module | FDR-corrected p-value group effect | FDR-corrected p-value time effect | FDR-corrected p-value group*time effect | Post-hoc analysis of significant group effect (in the absence of a significant group*time interaction): Dunnett's multiple comparisons with control test with sham/J60 group as control group: adjusted p-value | |
|---|---|---|---|---|---|
| Anterior DMLN | 0.0921 | 0.0150* | **0.0062** ** | | |
| Posterior DMN | <0.0001*** | <0.0001*** | **<0.0001** *** | | |
| Hippocampus | <0.0001*** | <0.0001*** | **<0.0001** *** | | |
| LCN | 0.5379 | 0.2247 | 0.5068 | | |
| SN | **0.0095** ** | 0.2174 | 0.1399 | sham/saline | 0.3284 |
| | | | | cholDREADD/J60 | 0.1546 |
| | | | | DREADD/J60 | 0.9569 |

*Supplementary Table 5. Statistical analysis of the ROI-based functional connectivity analysis of the pharmacological functional connectivity fMRI data with J60 or saline: post-hoc analysis per group of the significant group*time interactions from Supplementary Table 4. DMLN = default mode-like network, p.i. = post-injection. *p≤0.05, **p≤0.01, ***p≤0.001*

| **Network module** | **Post-hoc analysis of group*time interaction: analysis per group Time effects per group (FDR-corrected p-values)** | | **Post-hoc analysis of time effect: Dunnett's multiple comparisons with control test with baseline as control: adjusted p-value + direction of effect vs. baseline** |
|---|---|---|---|
| Anterior DMLN | sham/saline | 0.0699 | |
| | sham/J60 | 0.1827 | |
| | **cholDREADD/J60** | **0.0020**** | 10-20 min: **0.0026**(<)**, 20-30 min: **0.0041**(<)**, 30-40 min: **0.0206*(<)**, 40-50 min: 0.3934, 50-60 min p.i.: 0.9965 |
| | **DREADD/J60** | **0.0156*** | 10-20 min: **0.0375*(<)**, 20-30 min: 0.7295, 30-40 min: 1.0000, 40-50 min: 0.9998, 50-60 min p.i.: 0.6487 |
| Posterior DMLN | sham/saline | 0.1096 | |
| | sham/J60 | 0.0843 | |
| | **cholDREADD/J60** | **<0.0001***** | 10-20, 20-30, 30-40, 40-50, 50-60 min p.i.: **<0.0001***(<)** |
| | **DREADD/J60** | **<0.0001***** | 10-20, 20-30, 30-40, 40-50, 50-60 min p.i.: **<0.0001***(<)** |
| Hippocampus | sham/saline | 0.5571 | |
| | sham/J60 | 0.1058 | |
| | cholDREADD/J60 | 0.5049 | |
| | **DREADD/J60** | **<0.0001***** | 10-20 min: 0.2375, 20-30 min: 0.1401, 30-40 min: **0.0005***(>)**, 40-50, 50-60 min p.i.: **<0.0001***(>)** |

*Supplementary Table 6. Statistical analysis of the ROI-based functional connectivity analysis of the pharmacological functional connectivity fMRI data with J60 or saline: post-hoc analysis per time interval of the significant group*time interactions from Supplementary Table 4. DMLN = default mode-like network, p.i. = post-injection. *p≤0.05, **p≤0.01, ***p≤0.001*

| **Network module** | **Post-hoc analysis of group*time interaction: analysis per time interval Group effects per time interval (FDR-corrected p-values)** | | **Post-hoc analysis of group effect: Dunnett's multiple comparisons with control test with sham/J60 group as control group: adjusted p-value + direction of effect vs. control group** |
|---|---|---|---|
| Anterior DMLN | Baseline | 0.5989 | |
| | 10-20 min p.i. | **0.0024**** | sham/saline: 0.2906, cholDREADD/J60: **0.0006***(<)**, DREADD/J60: **0.0009***(<)** |
| | 20-30 min p.i. | **0.0114*** | sham/saline: 0.5822, cholDREADD/J60: **0.0016**(<)**, DREADD/J60: 0.0605 |
| | 30-40 min p.i. | 0.2378 | |
| | 40-50 min p.i. | 0.2378 | |
| | 50-60 min p.i. | 0.5989 | |
| Posterior DMLN | Baseline | 0.9830 | |
| | 10-20 min p.i. | **<0.0001***** | sham/saline: 0.8406, cholDREADD/J60: **<0.0001***(<)**, DREADD/J60: **0.0001***(<)** |
| | 20-30 min p.i. | **<0.0001***** | sham/saline: 0.9942, cholDREADD/J60: **<0.0001***(<)**, DREADD/J60: **0.0002***(<)** |
| | 30-40 min p.i. | **<0.0001***** | sham/saline: 0.9997, cholDREADD/J60: **0.0004***(<)**, DREADD/J60: **0.0004***(<)** |
| | 40-50 min p.i. | **<0.0001***** | sham/saline: 1.0000, cholDREADD/J60: **0.0039**(<)**, DREADD/J60: **0.0013**(<)** |
| | 50-60 min p.i. | **0.0006***** | sham/saline: 0.9833, cholDREADD/J60: **0.0154*(<)**, DREADD/J60: **0.0039**(<)** |
| Hippocampus | Baseline | 0.4810 | |
| | 10-20 min p.i. | **0.0147*** | sham/saline: 0.9103, cholDREADD/J60: **0.0492*(<)**, DREADD/J60: 0.8174 |
| | 20-30 min p.i. | **0.0245*** | sham/saline: 0.7838, cholDREADD/J60: **0.0494*(<)**, DREADD/J60: 0.9576 |
| | 30-40 min p.i. | **0.0028**** | sham/saline: 0.5317, cholDREADD/J60: 0.1289, DREADD/J60: 0.1796 |
| | 40-50 min p.i. | **<0.0001***** | sham/saline: 0.8522, cholDREADD/J60: 0.1110, DREADD/J60: **0.0031**(>)** |
| | 50-60 min p.i. | **<0.0001***** | sham/saline: 0.6165, cholDREADD/J60: 0.9996, DREADD/J60: **<0.0001*** (>)** |

*Supplementary Table 7. Statistical analysis of the seed-based functional connectivity analysis of the pharmacological functional connectivity fMRI data with J60 or saline: main effects and post-hoc analysis of the significant group and time effects. *p≤0.05, **p≤0.01, ***p≤0.001*

| Seed | FDR-corrected p-value group effect | FDR-corrected p-value time effect | FDR-corrected p-value group*time effect | Post-hoc analysis of significant group effect: Dunnett's multiple comparisons with control test with sham/J60 group as control group: adjusted p-value + direction of effect | | Post-hoc analysis of significant time effect: Dunnett's multiple comparisons with control test with baseline as control: adjusted p-value + direction of effect | |
|---|---|---|---|---|---|---|---|
| Orbitofrontal cortex | 0.0001*** | <0.0001*** | **<0.0001*** ** | | | | |
| Prelimbic-infralimbic cortex | **0.0159*** | **<0.0001*** ** | 0.3184 | sham/saline | 0.4156 | 10-20 min p.i. | **0.0005*** (<)** |
| | | | | cholDREADD/J60 | **0.0043** (<)** | 20-30 min | 0.4153 |
| | | | | | | 30-40 min | 0.9695 |
| | | | | DREADD/J60 | 0.0590 | 40-50 min | 0.2108 |
| | | | | | | 50-60 min | **0.0002*** (>)** |
| Anterior cingulate cortex | 0.0009*** | <0.0001*** | **0.0007*** ** | | | | |
| Posterior cingulate cortex | 0.0001*** | <0.0001*** | **<0.0001*** ** | | | | |
| Retrosplenial cortex | <0.0001*** | <0.0001*** | **<0.0001*** ** | | | | |
| Parietal association cortex | 0.0063** | 0.0002*** | **0.0007*** ** | | | | |
| Posterior parietal cortex | 0.0022** | <0.0001*** | **<0.0001*** ** | | | | |
| Anterior hippocampus | **0.0005*** ** | **0.0079*** ** | 0.2999 | sham/saline | 0.9921 | 10-20 min p.i. | 0.0535 |
| | | | | cholDREADD/J60 | **0.0010*** (<)** | 20-30 min | 0.2674 |
| | | | | | | 30-40 min | 0.9695 |
| | | | | DREADD/J60 | 0.9985 | 40-50 min | 0.7190 |
| | | | | | | 50-60 min | 0.9303 |
| Posterior hippocampus | <0.0001*** | <0.0001*** | **0.0024*** ** | | | | |
| Dentate gyrus | 0.0139* | 0.2205 | **0.0358*** | | | | |
| Nucleus accumbens | 0.2872 | 0.4186 | **0.0439*** | | | | |
| Caudate putamen | 0.0996 | 0.3746 | 0.3184 | | | | |
| Insular cortex | **0.0490*** | 0.4186 | 0.9854 | sham/saline | 0.9983 | | |
| | | | | cholDREADD/J60 | 0.0829 | | |
| | | | | DREADD/J60 | 0.9931 | | |

*Supplementary Table 8. Statistical analysis of the seed-based functional connectivity analysis of the pharmacological functional connectivity fMRI data with J60 or saline: post-hoc analysis per group of the significant group*time interactions from Supplementary Table 7. P.i. = post-injection. *p≤0.05, **p≤0.01, ***p≤0.001*

| **Seed** | **Post-hoc analysis of group*time interaction: analysis per group Time effects per group (FDR-corrected p-values)** | | **Post-hoc analysis of time effect: Dunnett's multiple comparisons with control test with baseline as control: adjusted p-value + direction of effect vs. baseline** |
|---|---|---|---|
| Orbitofrontal cortex | sham/saline | 0.6274 | |
| | sham/J60 | 0.1387 | |
| | cholDREADD/J60 | **<0.0001*****| 10-20, 20-30, 30-40, 40-50, 50-60 min p.i.: **<0.0001***(<)** |
| | DREADD/J60 | **0.0008***** | 10-20 min: <**0.0001***(<)**, 20-30 min: **0.0015**(<)**, 30-40 min and 40-50 min: **0.0016**(<)**, 50-60 min p.i.: **0.0114*(<)** |
| Anterior cingulate cortex | sham/saline | 0.5812 | |
| | sham/J60 | 0.5812 | |
| | cholDREADD/J60 | **<0.0001***** | 10-20, 20-30, 30-40 min: **<0.0001***(<)**, 40-50 min: **0.0018**(<)**, 50-60 min p.i.: 0.4756 |
| | DREADD/J60 | **0.0012**** | 10-20 min: **0.0004***(<)**, 20-30 min: **0.0235*(<)**, 30-40 min: 0.0627, 40-50 min: 0.1146, 50-60 min p.i.: 0.9999 |
| Posterior cingulate cortex | sham/saline | 0.3368 | |
| | sham/J60 | 0.3368 | |
| | cholDREADD/J60 | **<0.0001***** | 10-20, 20-30, 30-40, 40-50, 50-60 min p.i.: **<0.0001***(<)** |
| | DREADD/J60 | **<0.0001***** | 10-20, 20-30, 30-40, 40-50 min: **<0.0001***(<)**, 50-60 min p.i.: **0.0002***(<)** |
| Retrosplenial cortex | sham/saline | 0.5770 | |
| | sham/J60 | **0.0459*** | 10-20 min: 0.7963, 20-30 min: 0.3617, 30-40 min: 0.4022, 40-50 min: **0.0056**(<)**, 50-60 min p.i.: 0.1295 |
| | cholDREADD/J60 | **<0.0001***** | 10-20, 20-30, 30-40, 40-50, 50-60 min p.i.: **<0.0001***(<)** |
| | DREADD/J60 | **<0.0001***** | 10-20, 20-30, 30-40, 40-50, 50-60 min p.i.: **<0.0001***(<)** |
| Parietal association cortex | sham/saline | 0.2511 | |
| | sham/J60 | 0.1891 | |
| | cholDREADD/J60 | **<0.0001***** | 10-20, 20-30, 30-40, 40-50, 50-60 min p.i.: **<0.0001***(<)** |
| | DREADD/J60 | 0.1628 | |
| Posterior parietal cortex | sham/saline | 0.0839 | |
| | sham/J60 | 0.5809 | |
| | cholDREADD/J60 | **<0.0001***** | 10-20, 20-30, 30-40, 40-50, 50-60 min p.i.: **<0.0001***(<)** |
| | DREADD/J60 | 0.0638 | |
| Posterior hippocampus | sham/saline | 0.4941 | |
| | sham/J60 | 0.5117 | |
| | cholDREADD/J60 | **0.0001***** | 10-20 min: **<0.0001***(<)**, 20-30 min: **0.0002***(<)**, 30-40 min: **0.0113*(<)**, 40-50 min: **0.0014**(<)**, 50-60 min p.i.: **0.0366*(<)** |
| | DREADD/J60 | **0.0002***** | 10-20 min: 0.4533, 20-30 min: 0.6579, 30-40 min: 0.5644, 40-50 min: 0.1548, 50-60 min p.i.: **0.0223*(>)** |
| Dentate gyrus | sham/saline | 0.2660 | |
| | sham/J60 | 0.8963 | |
| | cholDREADD/J60 | **<0.0001***** | 10-20, 20-30, 30-40, 40-50 min: **<0.0001***(<)**, 50-60 min p.i.: **0.0196*(<)** |
| | DREADD/J60 | 0.9387 | |
| Nucleus accumbens | sham/saline | 0.5784 | |
| | sham/J60 | 0.8217 | |
| | cholDREADD/J60 | **0.0036**** | 10-20 min: **0.0017**(>)**, 20-30 min: **0.0015**(>)**, 30-40 min: **0.0057**(>)**, 40-50 min: **0.0009***(>)**, 50-60 min p.i.: 0.1710 |
| | DREADD/J60 | 0.6211 | |

*Supplementary Table 9. Statistical analysis of the seed-based functional connectivity analysis of the pharmacological functional connectivity fMRI data with J60 or saline: post-hoc analysis per time interval of the significant group*time interactions from Supplementary Table 7. P.i. = post-injection. *p≤0.05, **p≤0.01, ***p≤0.001*

| Seed | Post-hoc analysis of group*time interaction: analysis per time interval Group effects per time interval (FDR-corrected p-values) | | Post-hoc analysis of group effect: Dunnett's multiple comparisons with control test with sham/J60 group as control group: adjusted p-value + direction of effect vs. control group |
|---|---|---|---|
| Orbitofrontal cortex | Baseline | 0.5653 | |
| | 10-20 min p.i. | <0.0001*** | sham/saline: 0.1482, cholDREADD/J60: <0.0001***(<), DREADD/J60: <0.0001***(<) |
| | 20-30 min p.i. | <0.0001*** | sham/saline: 0.4221, cholDREADD/J60: <0.0001***(<), DREADD/J60: <0.0001***(<) |
| | 30-40 min p.i. | <0.0001*** | sham/saline: 0.8702, cholDREADD/J60: 0.0429*(<), DREADD/J60: 0.0099**(<) |
| | 40-50 min p.i. | 0.0008*** | sham/saline: 0.8196, cholDREADD/J60: 0.0396*(<), DREADD/J60: 0.0048**(<) |
| | 50-60 min p.i. | 0.0576 | |
| Anterior cingulate cortex | Baseline | 0.8165 | |
| | 10-20 min p.i. | <0.0001*** | sham/saline: 0.9234, cholDREADD/J60: <0.0001***(<), DREADD/J60: 0.0002***(<) |
| | 20-30 min p.i. | <0.0001*** | sham/saline: 0.9726, cholDREADD/J60: 0.0002***(<), DREADD/J60: 0.0012**(<) |
| | 30-40 min p.i. | 0.0004*** | sham/saline: 0.4208, cholDREADD/J60: 0.0002***(<), DREADD/J60: 0.0017**(<) |
| | 40-50 min p.i. | 0.0155* | sham/saline: 0.9903, cholDREADD/J60: 0.0548, DREADD/J60: 0.0447*(<) |
| | 50-60 min p.i. | 0.3644 | |
| Posterior cingulate cortex | Baseline | 0.9910 | |
| | 10-20 min p.i. | <0.0001*** | sham/saline: 0.5577, cholDREADD/J60: <0.0001***(<), DREADD/J60: <0.0001***(<) |
| | 20-30 min p.i. | <0.0001*** | sham/saline: 0.7972, cholDREADD/J60: <0.0001***(<), DREADD/J60: 0.0004***(<) |
| | 30-40 min p.i. | 0.0004*** | sham/saline: 0.9332, cholDREADD/J60: 0.0005***(<), DREADD/J60: 0.0060**(<) |
| | 40-50 min p.i. | 0.0005*** | sham/saline: 0.9372, cholDREADD/J60: 0.0026**(<), DREADD/J60: 0.0460*(<) |
| | 50-60 min p.i. | 0.0035** | sham/saline: 0.9927, cholDREADD/J60: 0.0126*(<), DREADD/J60: 0.0451*(<) |
| Retrosplenial cortex | Baseline | 0.5297 | |
| | 10-20 min p.i. | <0.0001*** | sham/saline: 0.3077, cholDREADD/J60: <0.0001***(<), DREADD/J60: <0.0001***(<) |
| | 20-30 min p.i. | <0.0001*** | sham/saline: 0.9993, cholDREADD/J60: <0.0001***(<), DREADD/J60: <0.0001***(<) |
| | 30-40 min p.i. | <0.0001*** | sham/saline: 0.8397, cholDREADD/J60: <0.0001***(<), DREADD/J60: <0.0001***(<) |
| | 40-50 min p.i. | <0.0001*** | sham/saline: 0.9811, cholDREADD/J60: 0.0007***(<), DREADD/J60: 0.0002***(<) |
| | 50-60 min p.i. | <0.0001*** | sham/saline: 1.0000, cholDREADD/J60: 0.0008***(<), DREADD/J60: <0.0001***(<) |
| Parietal association cortex | Baseline | 0.8567 | |
| | 10-20 min p.i. | 0.0048** | sham/saline: 0.9931, cholDREADD/J60: 0.0076**(<), DREADD/J60: 0.0756 |
| | 20-30 min p.i. | 0.0053** | sham/saline: 0.9754, cholDREADD/J60: 0.0239*(<), DREADD/J60: 0.0382*(<) |
| | 30-40 min p.i. | 0.0036** | sham/saline: 0.8268, cholDREADD/J60: 0.0012**(<), DREADD/J60: 0.0059**(<) |
| | 40-50 min p.i. | 0.0133* | sham/saline: 0.5279, cholDREADD/J60: 0.3408, DREADD/J60: 0.0958 |
| | 50-60 min p.i. | 0.0048** | sham/saline: 0.8016, cholDREADD/J60: 0.0280*(<), DREADD/J60: 0.0696 |
| Posterior parietal cortex | Baseline | 0.5831 | |
| | 10-20 min p.i. | 0.0003*** | sham/saline: 0.2802, cholDREADD/J60: 0.0002***(<), DREADD/J60: 0.0004***(<) |
| | 20-30 min p.i. | <0.0001*** | sham/saline: 0.9990, cholDREADD/J60: 0.0007***(<), DREADD/J60: 0.0035**(<) |
| | 30-40 min p.i. | 0.0004*** | sham/saline: 0.9929, cholDREADD/J60: 0.0022**(<), DREADD/J60: 0.0023**(<) |
| | 40-50 min p.i. | 0.0246* | sham/saline: 0.8672, cholDREADD/J60: 0.1881, DREADD/J60: 0.0814 |
| | 50-60 min p.i. | 0.0063** | sham/saline: 0.6724, cholDREADD/J60: 0.0557, DREADD/J60: 0.1215 |
| Posterior hippocampus | Baseline | 0.6455 | |
| | 10-20 min p.i. | 0.0003*** | sham/saline: 0.9935, cholDREADD/J60: 0.0004***(<), DREADD/J60: 0.1784 |
| | 20-30 min p.i. | 0.0003*** | sham/saline: 0.9753, cholDREADD/J60: 0.0002***(<), DREADD/J60: 0.0935 |
| | 30-40 min p.i. | 0.0006*** | sham/saline: 0.9891, cholDREADD/J60: 0.0014**(<), DREADD/J60: 0.8204 |
| | 40-50 min p.i. | <0.0001*** | sham/saline: 0.7640, cholDREADD/J60: 0.0002***(<), DREADD/J60: 0.9862 |
| | 50-60 min p.i. | 0.0003*** | sham/saline: 0.9684, cholDREADD/J60: 0.0167*(<), DREADD/J60: 0.2365 |
| Dentate gyrus | Baseline | 0.7784 | |
| | 10-20 min p.i. | 0.0267* | sham/saline: 0.9786, cholDREADD/J60: 0.0171*(<), DREADD/J60: 0.9994 |
| | 20-30 min p.i. | 0.0830 | |
| | 30-40 min p.i. | 0.0246* | sham/saline: 0.6614, cholDREADD/J60: 0.0035**(<), DREADD/J60: 0.8442 |
| | 40-50 min p.i. | 0.0658 | |
| | 50-60 min p.i. | 0.1850 | |
| Nucleus accumbens | Baseline | 0.3209 | |
| | 10-20 min p.i. | 0.1866 | |
| | 20-30 min p.i. | 0.3209 | |
| | 30-40 min p.i. | 0.3946 | |
| | 40-50 min p.i. | 0.3209 | |
| | 50-60 min p.i. | 0.5760 | |

*Supplementary Table 10. Statistical analysis of the ROI-based and seed-based functional connectivity analysis of the resting-state fMRI data of the memory experiment (pre-learning, post-acquisition training and post-reversal training): main effects. DMLN = default mode-like network, LCN = lateral cortical network, SN = salience-like network.*

| Network module or seed | FDR-corrected p-value group effect | FDR-corrected p-value time effect | FDR-corrected p-value group*time effect |
|---|---|---|---|
| Anterior DMLN | 0.7916 | 0.9727 | 0.7644 |
| Posterior DMLN | 0.7916 | 0.6170 | 0.3613 |
| Hippocampus | 0.7916 | 0.9727 | 0.7644 |
| LCN | 0.8175 | 0.9727 | 0.3613 |
| SN | 0.7916 | 0.9727 | 0.7912 |
| Orbitofrontal cortex | 0.8431 | 0.9316 | 0.8427 |
| Prelimbic-infralimbic cortex | 0.8431 | 0.6760 | 0.8889 |
| Anterior cingulate cortex | 0.8431 | 0.8896 | 0.8889 |
| Posterior cingulate cortex | 0.8431 | 0.6432 | 0.8889 |
| Retrosplenial cortex | 0.8431 | 0.6432 | 0.3393 |
| Parietal association cortex | 0.8431 | 0.6432 | 0.3393 |
| Posterior parietal cortex | 0.8431 | 0.9316 | 0.3598 |
| Anterior hippocampus | 0.8431 | 0.6432 | 0.1612 |
| Posterior hippocampus | 0.8431 | 0.6760 | 0.8889 |
| Dentate gyrus | 0.8431 | 0.6760 | 0.8889 |
| Nucleus accumbens | 0.8431 | 0.6760 | 0.8889 |
| Caudate putamen | 0.8431 | 0.9316 | 0.8973 |
| Insular cortex | 0.8431 | 0.6760 | 0.8889 |

*Supplementary Table 11. Statistical analysis of the ROI-based and seed-based functional connectivity analysis of the resting-state fMRI data of the food restriction (FR) experiment: main effect of time and post-hoc analysis of the significant time effects: comparison with pre-FR baseline and with 7-day (7d) FR timepoint. 28d no FR = 28 days no FR, after ending the 28d FR period. *p≤0.05, **p≤0.01, ***p≤0.001*

| **Network module or seed** | **FDR-corrected p-value time effect** | **Post-hoc analysis of time effect: Dunnett's multiple comparisons with control test with pre-FR baseline as control: adjusted p-value + direction of effect vs. control** | **Post-hoc analysis of time effect: Dunnett's multiple comparisons with control test with 7d FR as control: adjusted p-value + direction of effect vs. control** |
|---|---|---|---|
| Anterior DMLN | 0.0515 | 7d FR: 0.1931, 21d FR: 0.1367, 28d FR: **0.0037**(>)**, 28d no FR: 0.0705 | Pre-FR: 0.1886, 21d FR: 0.9998, 28d FR: 0.3536, 28d no FR: 0.9787 |
| Posterior DMLN | 0.0515 | 7d FR: 0.2894, 21d FR: 0.0779, 28d FR: **0.0031**(>)**, 28d no FR: 0.3849 | Pre-FR: 0.2830, 21d FR: 0.9264, 28d FR: 0.2146, 28d no FR: 0.9983 |
| Hippocampus | 0.1790 | | |
| LCN | 0.0805 | | |
| SN | 0.0805 | | |
| Orbitofrontal cortex | **0.0004*** ** | 7d FR: 0.0583, 21d FR: **0.0405*(>)**, 28d FR: **<0.0001***(>)**, 28d no FR: **0.0084**(>)** | Pre-FR: 0.0571, 21d FR: 1.0000, 28d FR: 0.0606, 28d no FR: 0.8950 |
| Prelimbic-infralimbic cortex | **0.0002*** ** | 7d FR: 0.2018, 21d FR: 0.1578, 28d FR: **<0.0001***(>)**, 28d no FR: **0.0126*(>)** | Pre-FR: 0.1973, 21d FR: 1.0000, 28d FR: **0.0049**(>)**, 28d no FR: 0.6128 |
| Anterior cingulate cortex | **0.0002*** ** | 7d FR: 0.0858, 21d FR: **0.0133*(>)**, 28d FR: **<0.0001***(>)**, 28d no FR: **0.0110*(>)** | Pre-FR: 0.0840, 21d FR: 0.8929, 28d FR: **0.0073**(>)**, 28d no FR: 0.8571 |
| Posterior cingulate cortex | **0.0003*** ** | 7d FR: 0.3367, 21d FR: **0.0121*(>)**, 28d FR: **<0.0001***(>)**, 28d no FR: **0.0179*(>)** | Pre-FR: 0.3292, 21d FR: 0.4080, 28d FR: **0.0050**(>)**, 28d no FR: 0.5006 |
| Retrosplenial cortex | **0.0003*** ** | 7d FR: 0.1275, 21d FR: **0.0064**(>)**, 28d FR: **<0.0001***(>)**, 28d no FR: 0.1078 | Pre-FR: 0.1248, 21d FR: 0.6066, 28d FR: **0.0237*(>)**, 28d no FR: 1.0000 |
| Parietal association cortex | **0.0002*** ** | 7d FR: 0.6004, 21d FR: 0.1371, 28d FR: **<0.0001***(>)**, 28d no FR: **0.0297*(>)** | Pre-FR: 0.5900, 21d FR: 0.7995, 28d FR: **0.0007***(>)**, 28d no FR: 0.3636 |
| Posterior parietal cortex | **<0.0001*** ** | 7d FR: 0.1262, 21d FR: **0.0053**(>)**, 28d FR: **<0.0001***(>)**, 28d no FR: 0.0740 | Pre-FR: 0.1235, 21d FR: 0.5636, 28d FR: **0.0006***(>)**, 28d no FR: 0.9990 |
| Anterior hippocampus | 0.1814 | | |
| Posterior hippocampus | **0.0002*** ** | 7d FR: 0.4129, 21d FR: 0.9770, 28d FR: **0.0002***(>)**, 28d no FR: 0.1772 | Pre-FR: 0.4042, 21d FR: 0.1963, 28d FR: **0.0160*(>)**, 28d no FR: 0.9746 |
| Dentate gyrus | **<0.0001*** ** | 7d FR: 0.6743, 21d FR: 0.9993, 28d FR: **<0.0001***(>)**, 28d no FR: **0.0047**(>)** | Pre-FR: 0.6648, 21d FR: 0.7806, 28d FR: **0.0012**(>)**, 28d no FR: 0.0825 |
| Nucleus accumbens | **0.0014** ** | 7d FR: 0.2801, 21d FR: 0.4647, 28d FR: **0.0002***(>)**, 28d no FR: 0.2959 | Pre-FR: 0.2739, 21d FR: 0.9882, 28d FR: **0.0325*(>)**, 28d no FR: 1.0000 |
| Caudate putamen | **0.0003*** ** | 7d FR: 0.7681, 21d FR: 0.0762, 28d FR: **<0.0001***(>)**, 28d no FR: **0.0083**(>)** | Pre-FR: 0.7594, 21d FR: 0.4497, 28d FR: **0.0018**(>)**, 28d no FR: 0.0948 |
| Insular cortex | **0.0012** ** | 7d FR: 0.2968, 21d FR: 0.3360, 28d FR: **0.0002***(>)**, 28d no FR: **0.0396*(>)** | Pre-FR: 0.2901, 21d FR: 0.9999, 28d FR: **0.0267*(>)**, 28d no FR: 0.7658 |
